# Menthol in Electronic Cigarettes: A Contributor to Respiratory Disease?

**DOI:** 10.1101/2020.03.14.988006

**Authors:** Vijayalekshmi Nair, Malcolm Tran, Rachel Z. Behar, Song Zhai, Xinping Cui, Rattapol Phandthong, Yuhuan Wang, Songqin Pan, Wentai Luo, James F. Pankow, David C. Volz, Prue Talbot

**Author notes:** Lead Contact and Corresponding Author: Prue Talbot. The first two authors contributed equally to this work.

## Abstract

Menthol is widely used in tobacco products. This study compared the effects of menthol on human bronchial epithelium using submerged cultures, a VITROCELL® cloud chamber that provides air liquid interface (ALI) exposure without solvents or heating, and a Cultex ALI system that delivers aerosol equivalent to that inhaled during vaping. In submerged culture, menthol significantly increased calcium influx and mitochondrial reactive oxygen species (ROS) via the TRPM8 receptor, responses that were inhibited by a TRPM8 antagonist. VITROCELL® cloud chamber exposure of BEAS-2B monolayers increased mitochondrial protein oxidation, expression of the antioxidant enzyme SOD2, activation of NF-κB, and secretion of inflammatory cytokines (IL-6 and IL-8). Proteomics data collected following ALI exposure of 3D EpiAirway tissue in the Cultex showed upregulation of NRF-2-mediated oxidative stress, oxidative phosphorylation, and IL-8 signaling. Across the three platforms, menthol adversely effected human bronchial epithelium in a manner that could lead to respiratory disease.

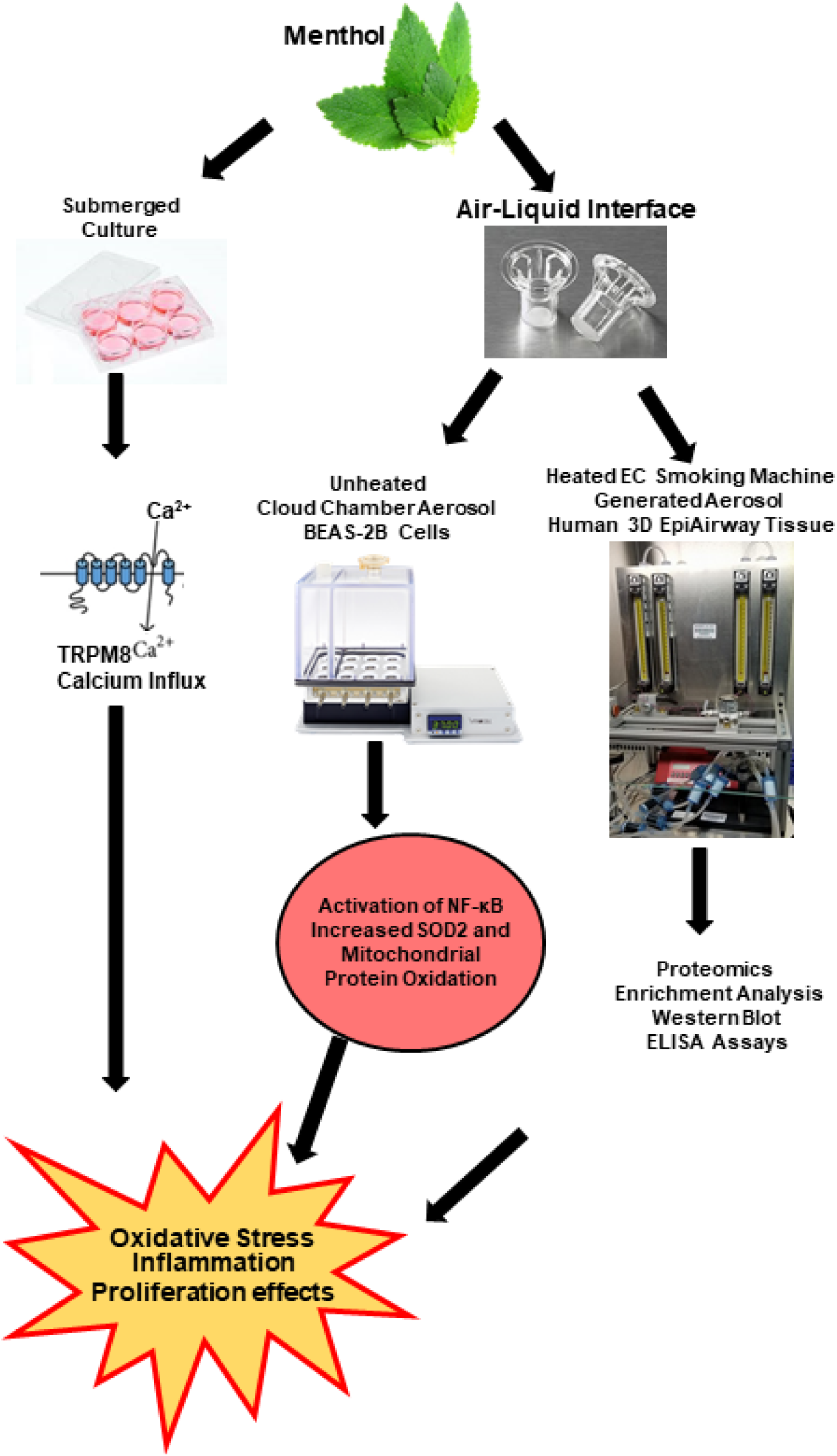
Graphical Abstract. Mechanism of action of menthol on human bronchial epithelium. Three *in vitro* platforms were used to study the effect of menthol on bronchial epithelium. In submerged culture (using BEAS-2B cells), menthol produced rapid calcium influx followed by an increase in oxidative stress and inflammatory cytokines. ALI exposure of BEAS-2B cells to unheated menthol in a cloud chamber caused activation of an inflammatory transcription factor (NF-κB) and oxidative stress. Proteomics analysis of human EpiAirway tissues exposed at the ALI to heated menthol EC aerosols identified changes in the expression of proteins involved in oxidative stress and in an inflammatory response.

## INTRODUCTION

Flavor chemicals are used in almost all tobacco products, including electronic cigarettes (ECs) (Behar et al., 2018; Hua et al., 2019; Lisko et al., 2014; Tierney et al., 2016), and numerous attractive flavors have contributed to the rapid rise in the popularity of ECs among all age groups in the US (U.S. Department of Health and Human Services, 2016; Miech et al., 2019; U.S. Department of Health and Human Services, 2016). While most flavor chemicals in consumer products are “generally regarded as safe” (GRAS), the Flavor and Extracts Manufacturers Association (FEMA) has cautioned that their GRAS designation pertains only to ingestion, not inhalation (Hallagan, 2014). Because the data on flavor chemical ingestion cannot be directly translated to inhalation, the health consequences of short-and long-term inhalation of flavor chemicals in ECs remain largely uncharacterized. This problem is compounded by the lack of validated methods for determining the effects of EC flavor chemicals and their reaction products on the respiratory system.

Menthol is often used in ECs (Behar et al., 2018; Hua et al., 2019) and is the only flavor chemical permitted in tobacco cigarettes under the Family Smoking Prevention and Tobacco Control Act (2009). EC refill fluids and conventional cigarettes sometimes contain menthol, even when they are sold as non-mentholated (Behar et al., 2018; Henderson, 2019; Omaiye et al., 2018). Menthol produces a cooling effect upon binding to the TRPM8 receptor (Transient Receptor Potential Melastatin 8), a cation channel with selectivity for calcium (Peier et al., 2002). Menthol is used in tobacco products to impart flavor and to reduce the harshness of inhaled tobacco smoke, making inhalation of tobacco aerosols easier for novices (DeVito et al., 2019; Willis et al., 2011).

Mentholated ECs may facilitate the initiation of smoking, increase nicotine dependence, and increase progression to conventional cigarette smoking (Food and Drug Administration, 2011; Nonnemaker et al., 2013; Villanti et al., 2017). Mentholated tobacco cigarettes also reduce cessation rates when compared to non-mentholated tobacco cigarettes (Delnevo et al., 2011). Mentholated tobacco cigarettes are widely distributed among the African American community and adolescent smokers, and are used more often by women than men (Food and Drug Administration, 2011). In a weight of evidence analysis on conventional cigarettes, it was concluded that menthol is not associated with a disease risk to the user (Food and Drug Administration, 2011). However, this conclusion was based on comparisons of mentholated and non-mentholated conventional cigarettes, and it may not pertain to ECs, which often have much higher concentrations of menthol than those in food and other consumer products, including tobacco cigarettes (Hua et al., 2019; Tierney et al., 2016). As examples, in mentholated tobacco cigarettes the concentration of menthol was reported in the range of 0.52-4.19 mg/cigarette (Ai et al., 2016), and in a second study, the average concentration was 4.75 mg/cigarette (Paschke et al., 2017). In contrast, menthol concentration in one EC refill fluid was 85 mg/mL (Behar et al., 2017) and 15 mg/mL in mint flavored JUUL pods (Omaiye et al., 2018), which are popular with high school-aged users (Barrington-Trimis and Leventhal, 2018).

Existing studies indicate a need for further work on the potential for high menthol concentrations in ECs to be associated with disease. For example, in submerged 2-dimensional (2D) cell cultures, EC refill fluids and aerosols had cytotoxic effects on adult and embryonic cells, and these were often associated with flavor chemical concentrations (Bahl et al., 2012; Behar et al., 2017; Hua et al., 2019). Pure menthol was cytotoxic to bronchial epithelium at the concentrations found in EC products when tested *in vitro* with the MTT assay using 2D submerged cell cultures (Behar et al., 2017; Hua et al., 2019). Lin et al., (2018) showed that subchronic exposure of mice to mentholated cigarette smoke induced more inflammation in lungs than smoke from non-mentholated cigarettes. Recently, serious respiratory illness and death have been attributed to EC use, and patients requiring hospitalization have been reported to have “e-cigarette or vaping product use-associated lung injury” (EVALI) (Centers for Disease Control and Prevention, 2019). The etiology of EVALI is not yet understood, but EC products with high concentrations of flavor chemicals should be investigated as possible causative agents.

The purpose of the current study was to understand how menthol, at the concentrations found in ECs, affects human respiratory epithelium and to compare responses to menthol across three *in vitro* platforms. In all protocols, the concentrations tested produced no effect in the MTT assay (referred to as the MTT NOAEL – no observed adverse effect level). In the first protocol, BEAS-2B cells from human bronchial epithelium were exposed to various concentrations of pure menthol using submerged 2D cultures and endpoints relating to oxidative stress and cell proliferation were examined. In the second approach, BEAS-2B cells were exposed at the air liquid interface (ALI) to menthol aerosols using a cloud chamber, which creates aerosol without heating or solvent (propylene glycol or PG) exposure. This was done to mimic *in vivo* exposure, while avoiding the possibility of producing heat-induced reaction products during aerosolization. Endpoints related to oxidative stress and cytokine signaling were examined. In the third protocol, 3D models of human respiratory epithelium (EpiAirway tissues) were exposed at the ALI to aerosol created by heating e-fluid in an EC using a smoking machine. This exposure was similar to actual vaping, as the heated aerosol contained menthol, PG, and any reaction products that formed during heating. Proteomics analysis was performed on the EpiAirway tissue exposed to menthol aerosol. Data were compared across the three *in vitro* platforms and evaluated for their potential to contribute to respiratory diseases, such as chronic obstructive pulmonary disease (COPD), emphysema, and EVALI. To give relevance to our data in the context of ECs, all menthol concentrations that we tested were within the range found in EC products (Behar et al., 2017; Hua et al., 2019), and sublethal concentrations were used in the three in vitro protocols.

## RESULTS

### Expression of TRPM8 Receptor

Menthol mediates signal transduction through the TRPM8 receptor, a ligand-gated cation channel with moderate to high selectivity for calcium ions (Peier et al., 2002). The expression of the TRPM8 receptor in human lung epithelial cells and lung fibroblasts was evaluated using western blotting and immunofluorescence microscopy (Figures 1A-C). Immunoreactivity of the TRPM8 receptor in BEAS-2B cells was intermediate between A549 cancer cells and human pulmonary fibroblasts (hPFs) (Figure 1A). The pattern of fluorescence was punctate and consistent with localization in the plasma membrane (Figure 1B). BEAS-2B cells treated with secondary antibody alone (negative control) had no label (Figure 1C).

**Figure 1.**
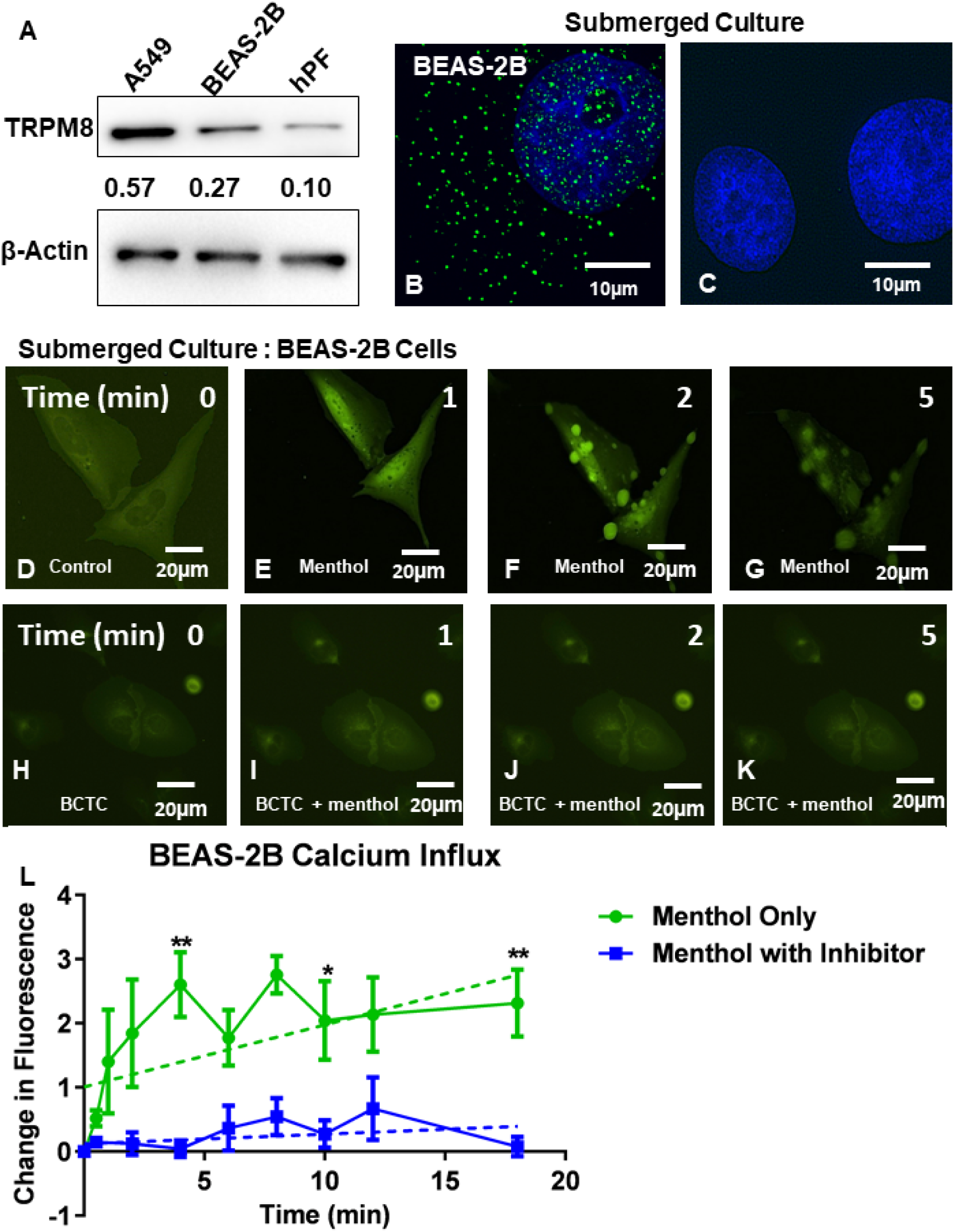
Menthol Induces Calcium Influx via the TRPM8 Receptor in Submerged Cultures of BEAS-2B Cells. **(A)** TRPM8 western blot of A549 cells, BEAS-2B cells, and hPFs with β-actin as the loading control. **(B-C)** Immunocytochemical staining of BEAS-2B cells with a human TRPM8 antibody (B), and negative control treated with secondary antibody alone (C). The nuclei were counterstained using DAPI. This experiment was performed three times. **(D-K)** Time-lapse micrographs of BEAS-2B cells transfected with the GCaMP5 plasmid and treated with 0.2 mg/mL (1.3 mM) of pure menthol (D-G) and TRPM8 inhibitor (BCTC) plus menthol (H-K). This experiment was performed three times. **(L)** Graph showing changes in fluorescence intensity in menthol-treated cells with and without the TRPM8 inhibitor. A two-way ANOVA was performed by comparing change in green fluorescence versus time, and significant changes in green fluorescence are indicated by ** and * for p <0.01 and p <0.05, respectively. Each point is the mean of three experiments ± the standard error of the mean (SEM).

### Menthol Fluids and Aerosol Fluids Were Cytotoxic in Submerged Cultures

The cytotoxicity of pure menthol in culture medium (menthol fluid) and menthol aerosols dissolved in medium (hereafter referred to as aerosol fluid) were examined in submerged cultures using the MTT assay (Supplementary Figures 1A, B). Test solutions were considered cytotoxic if absorbance was reduced to < IC_70_ (reduction of 30% relative to the untreated control) according ISO protocol #10993-5 (ISO-10993-5-2009). Menthol fluids were cytotoxic in a concentration-dependent manner with the IC_70_ and IC_50_ values being 0.26 mg/mL and 0.87 mg/mL, respectively (Supplementary Figure 1A). Menthol concentrations as low as 0.93 mg/mL caused a significant reduction relative to the control (p <0.01) in the fluid group. Menthol aerosol fluids were likewise cytotoxic producing an IC_70_ at 0.369 mg/mL. (Supplementary Figure 1B).

### In Submerged Cultures Menthol Induced Calcium Influx in BEAS-2B Cells through Activation of TRPM8 Receptor

The effect of menthol on calcium influx was measured in BEAS-2B cells using GCaMP5, a genetically encoded calcium indicator plasmid (Ackerboom et al., 2012). BEAS-2B cells transfected with GCaMP5 were treated with 0.2 mg/mL of menthol (MTT NOAEL) and time-lapse video was collected (Figures 1D-G). Intracellular fluorescence was low prior to treatment (Figure 1D). There was a rapid increase in cytosolic calcium indicated by increased green fluorescence during the first minute of menthol treatment. Calcium was initially high in the perinuclear region (Figure 1E) and later became concentrated in large vesicles that were highly fluorescent (Figure 1F, Supplementary Video 1). These vesicles bulged from the surface of the cells but were not exocytosed. Pretreatment of cells with 10 µM BCTC (an antagonist of the TRPM8 receptor) prior to menthol treatment attenuated calcium influx caused by menthol (Figures. 1 H-K). The time-lapse data were quantified, and significant differences were seen between the menthol treated group and the group pre-incubated with BCTC prior to menthol treatment (Figure 1L). These data indicate that menthol caused calcium influx by activation of the TRPM8 receptor and not non-specific disruption of the cell plasma membrane.

### Menthol Treatment Inhibited Cell Proliferation in Submerged Cultures

Live cell imaging and video bioinformatics software were used to investigate the effect of menthol fluid and menthol aerosol fluid on cell morphology, proliferation, and survival (Figure 2). BEAS-2B cells were treated with either 0.02 mg/mL (low) or 0.2 mg/mL (high) concentrations of menthol fluid or menthol aerosol fluid. Treatments were done either during plating of cells (attaching) (Figures 2 A-J) or after cells had been plated and attached for 24 h (Figures 2K-T).

**Figure 2.**
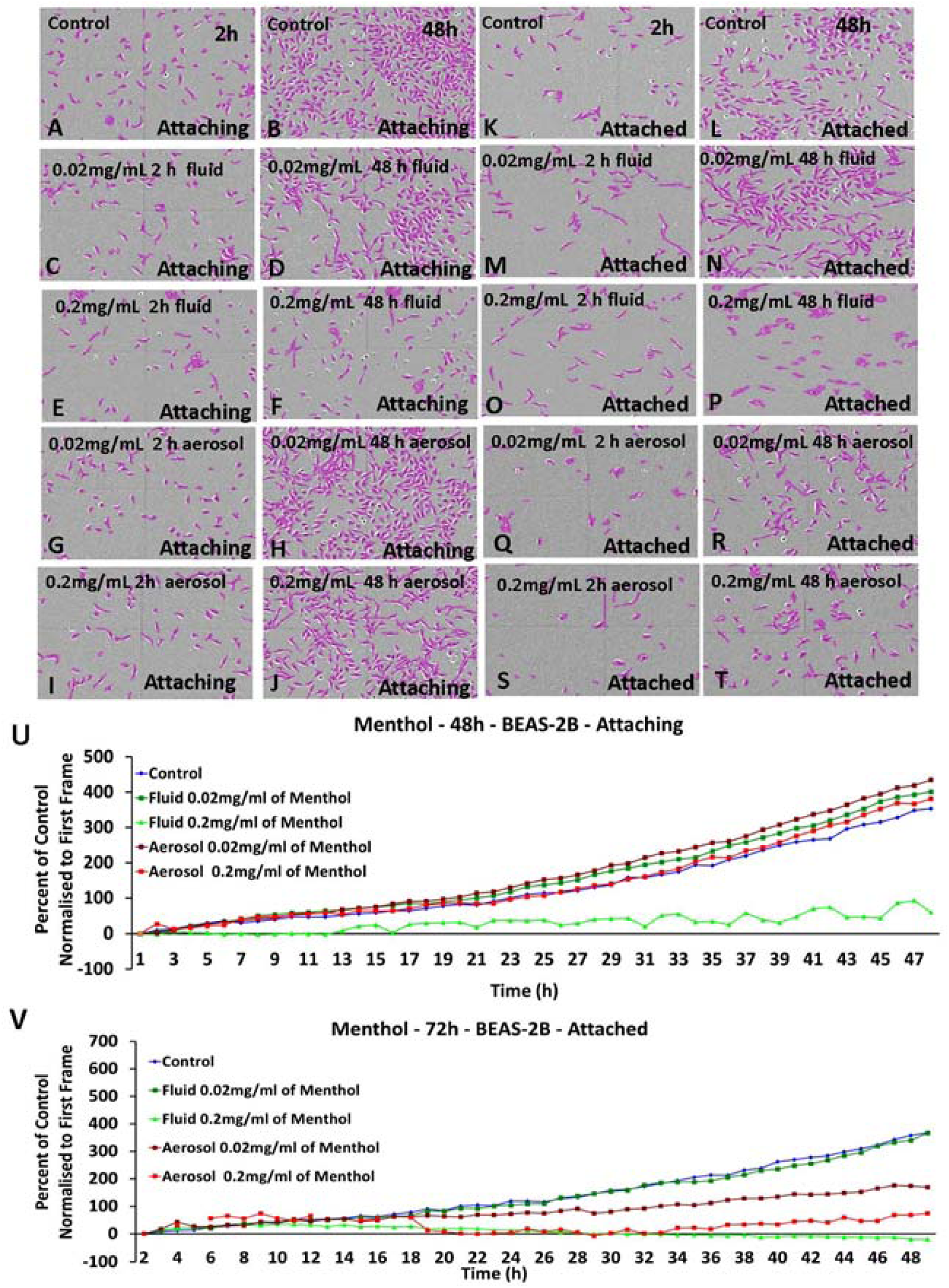
Effect of Menthol Fluids and Aerosol Fluids on Proliferation of BEAS-2B Cells in Submerged Culture. **(A-T)** Micrographs of BEAS-2B cells treated with menthol fluid (0.02 mg/mL or 0.2 mg/mL) and menthol aerosol fluid (0.02 mg/mL and 0.2 mg/mL) during plating (A-J attaching) and 24 h after plating (K-T attached). Cells were imaged live in a Nikon BioStation CT, and time-lapse images were captured every 2 h for 48 h. Cells have been segmented with CL-Quant software and colorized to show their boundaries clearly. **(U and V)** Graphs showing confluency of treated cells normalized to untreated controls versus time in control and treatment groups. Data are plotted as means of 2 experiments.

The low concentration of both menthol fluid and aerosol fluid did not affect proliferation of attaching cells (Figures 2D, H, and U). However, when attaching BEAS-2B cells were treated with the high concentration of menthol fluid during plating, they did not proliferate (Figure 2F). In contrast, the high concentration (0.2 mg/mL) of menthol aerosol fluid did not affect proliferation of attaching cells (Figure 2J), probably because not all menthol transferred to the aerosol fluid.

When attached BEAS-2B cells were treated with the high concentration (0.2 mg/mL) of menthol fluid or menthol aerosol fluid, proliferation was significantly decreased (Figures 2P,T and V), while the low concentration of aerosol fluid had an effect intermediate between the high concentration and the untreated controls (Figures 2R and V). The low concentration of fluid did not affect attached cells (Figure 2N).

### ROS Generation in Submerged Cultures

To investigate effects downstream of menthol-induced calcium elevation, intracellular ROS was measured in menthol fluid-exposed BEAS-2B cells in submerged culture. Superoxide (O2•-) generated from mitochondrial oxidative phosphorylation is a major source of cellular ROS. MitoSOX Red, a fluorescent indicator specific for superoxide, was used to localize and quantify superoxide in menthol-treated cells. Live cell imaging results for BEAS-2B cells incubated with MitoSOX showed increased mitochondrial ROS generation in menthol fluid (0.2 mg/mL, MTT NOAEL) treated cells (Figures. 3A, B). Menthol induced mitochondrial ROS was decreased when cells were pretreated with BCTC prior to menthol exposure (Figures. 3C, D)

**Figure 3.**
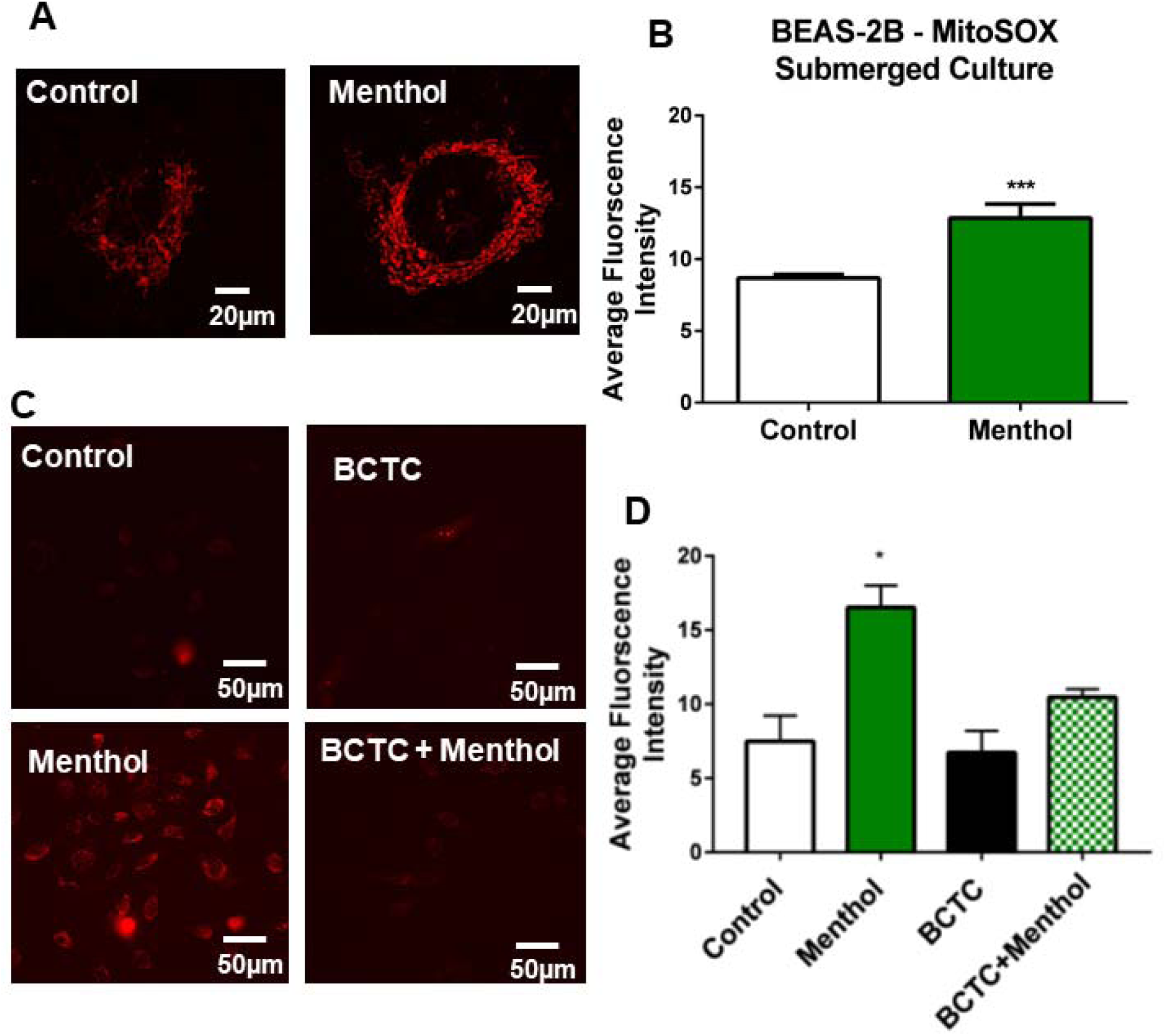
Mitochondrial ROS Generation in Menthol-treated BEAS-2B Cells in Submerged Culture. **(A)** Micrographs of BEAS-2B cells labeled with MitoSOX-Red after no treatment (control) or treatment with 0.2 mg/mL menthol. **(B)** Graph showing average fluorescence intensity per cell in control and menthol-treated cells. A two tailed t-test was used to compare fluorescent intensity. In B, each bar is the mean of three experiments ± the standard error of the mean (SEM). *** = p<0.0001 **(C and D)** Effects of menthol on BEAS-2B cells after blocking the TRPM8 receptor with BCTC. Cells were labeled with MitoSOX-Red after menthol treatment (4 h) with and without TRPM8 inhibitor (BCTC). Statistical significance was determined using a one-way ANOVA and significant changes were isolated using Dunnett’s posthoc test in which each group was compared to the untreated control. In D, each bar is the mean of three experiments ± the standard error of the mean. * = p < 0.05; *** = p<0.00.

### Oxidative Stress Occurs During ALI Exposure of BEAS-2B Monolayers to Unheated Menthol Aerosol Generated Using a Cloud Chamber

The preceding studies were done using submerged cultures. A VITROCELL® cloud chamber was used to determine how monolayers of BEAS-2B cells respond when exposed to menthol at the air-liquid interface (ALI). The cloud chamber creates an aerosol without heating, without use of a solvent such as PG, and without introduction of heat-induced reaction products. Menthol (0.8 mg/mL) aerosol was generated in the cloud chamber as described in the Transparent Methods. The actual concentration of menthol in the aerosol in both the VITROCELL® and Cultex experiments was not directly measured, but 100% transfer was assumed. The MTT assay indicated that cytotoxicity (absorbance < 70% of the control) was not induced by menthol in BEAS-2B cells using our exposure protocol in the ALI VITROCELL® cloud chamber (Figure 4A).

**Figure 4.**
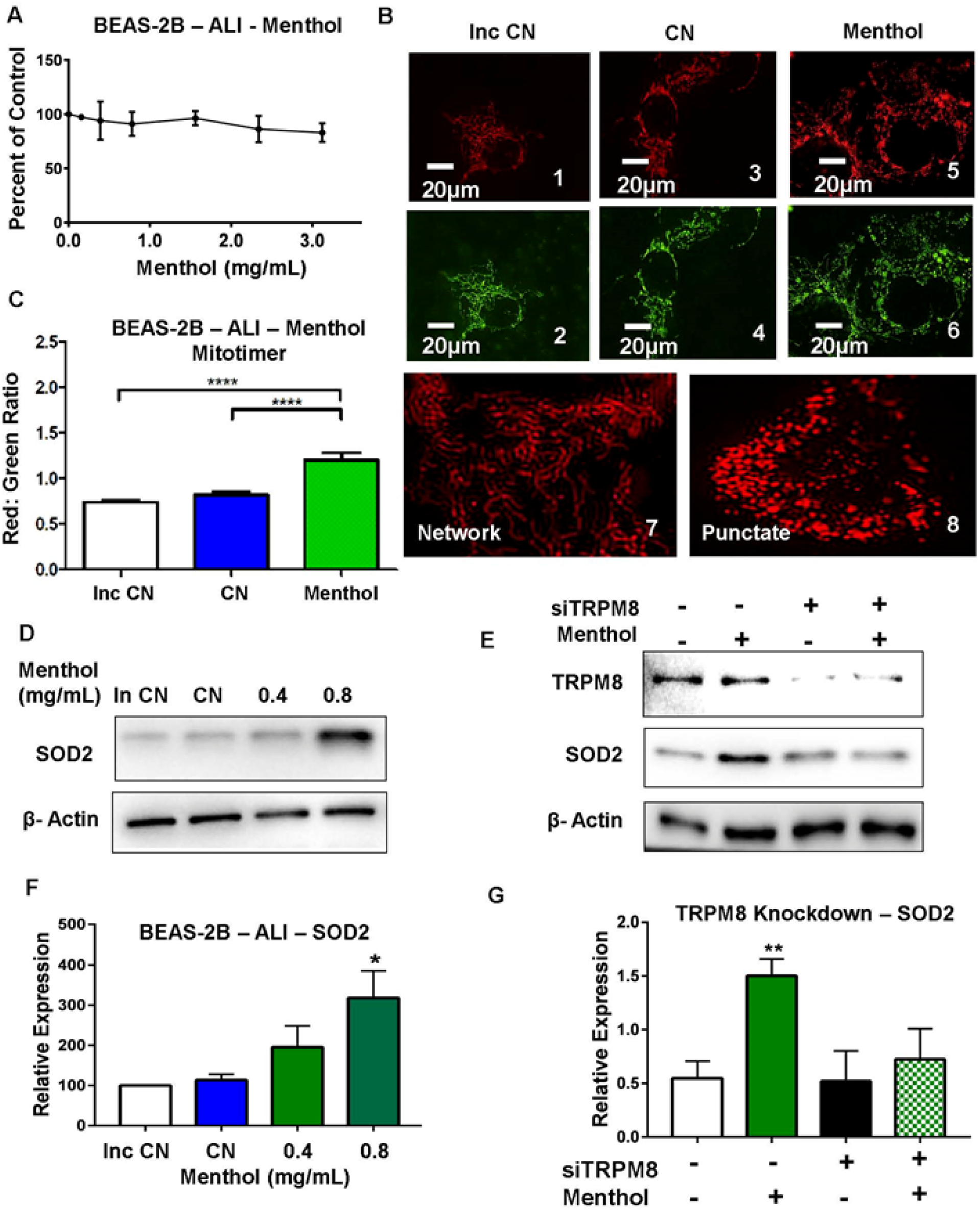
Menthol Exposure at the ALI in a Cloud Chamber Induced an Oxidative Stress Response and Elevation of an Antioxidant Enzyme. **(A)** MTT dose-response curve showing absorbance (percent of control) plotted as a function of different concentrations (0.15 – 3.125 mg/mL) of menthol aerosol in ALI exposure. Monolayers of BEAS-2B cells were used in all experiments. **(B)** Fluorescent micrographs of BEAS-2B cells transfected with MitoTimer plasmid. (1, 2) are micrographs of the incubator control, (3, 4) are control aerosol exposure and (5, 6) are menthol aerosol exposure. (7, 8) are magnified images showing networked and punctate mitochondria before and after menthol treatment. **(C)** The red/green ratio of the MitoTimer expressing cells is plotted for each group. Each bar is the mean of three experiments ± the standard error of the mean. A one-way ANOVA was used to compare means. **** = p< 0.00001 **(D)** Expression of SOD2 in BEAS-2B cells exposed to menthol aerosol (0.4 β-actin was used as the loading control. Inc CN is the incubator control, and CN is the control exposed to 1% DMSO. **(E)** BEAS-2B cells were treated with siRNA against TRPM8 and exposed to menthol aerosol. Whole cell lysates were then analyzed by western blot for β-actin was used as the loading control **(F and G)** Relative expression of SOD2 in western blots D and E respectively. Bars in F and G are means of three independent experiments and error bars represent the standard error of the means. A one-way ANOVA with Dunnett’s posthoc test was used to compare means in the knockdown experiment. * = p< 0.05, ** = p<0.01.

To visualize mitochondria and oxidation of mitochondrial proteins, we transfected cells with the MitoTimer plasmid, which is targeted to mitochondria via cytochrome c (Laker et al., 2014). MitoTimer fluoresces green when mitochondrial protein is not oxidized. As protein oxidation increases, the fluorescence shifts from green to red. Cells transfected with the MitoTimer plasmid were exposed to menthol aerosol (0.8 mg/mL) in the VITROCELL® cloud chamber as described in the Materials and Methods. Ratiometric analysis of red/green MitoTimer fluorescence revealed a statistically significant increase of mitochondrial protein oxidation in menthol treated cells (Figures 4B, C). A change in mitochondrial morphology was also observed in treated cells (Figure 4B). Mitochondria were predominantly networked in control cells (Figure 4B1, *2, 3,* 4) and punctate after menthol treatment (Figure 4B: Micrographs 5, 6, 7, and 8)

Cellular ROS levels are regulated by antioxidant systems. The most crucial antioxidant is manganese superoxide dismutase (MnSOD/SOD-2), which neutralizes superoxide by converting it into hydrogen peroxide (H_2_O_2_) (Holley et al., 2011). Aerosol generated using 0.8 mg/mL of menthol increased expression of SOD2 in a concentration dependent manner, as shown in western blots (Figures 4D, F). We next investigated the effect of TRPM8 silencing on SOD2 expression. Knockdown of TRPM8 using siRNA prior to menthol exposure significantly reversed the effect of menthol aerosol on SOD2 levels in treated cells (Figures 4 E, G).

### Activation of Nuclear Factor Kappa B (NF-κB) is Stimulated by ALI Exposure to Unheated Menthol Aerosol Generated Using a Cloud Chamber

NF-κB is a transcription factor that is activated in response to several stimuli, including oxidative stress. To evaluate the role of menthol in NF-κB activation, cells were exposed to menthol aerosol (0.8 mg/mL) in a VITROCELL® cloud chamber as described in the Transparent Methods. 24 h after menthol aerosol exposure, there was a significant increase in phospho-NF-κB (active form) expression in the whole cell lysate when compared to the untreated control (Figure 5A). To assess the translocation of phospho-NF-κB into the nucleus, cells treated with or without menthol aerosol were subjected to cell fractionation to separate nuclear and cytoplasmic proteins. There was a significant increase in phospho-NF-κB in the nuclear fraction of cells exposed to menthol aerosol (Figure 5B).

**Figure 5.**
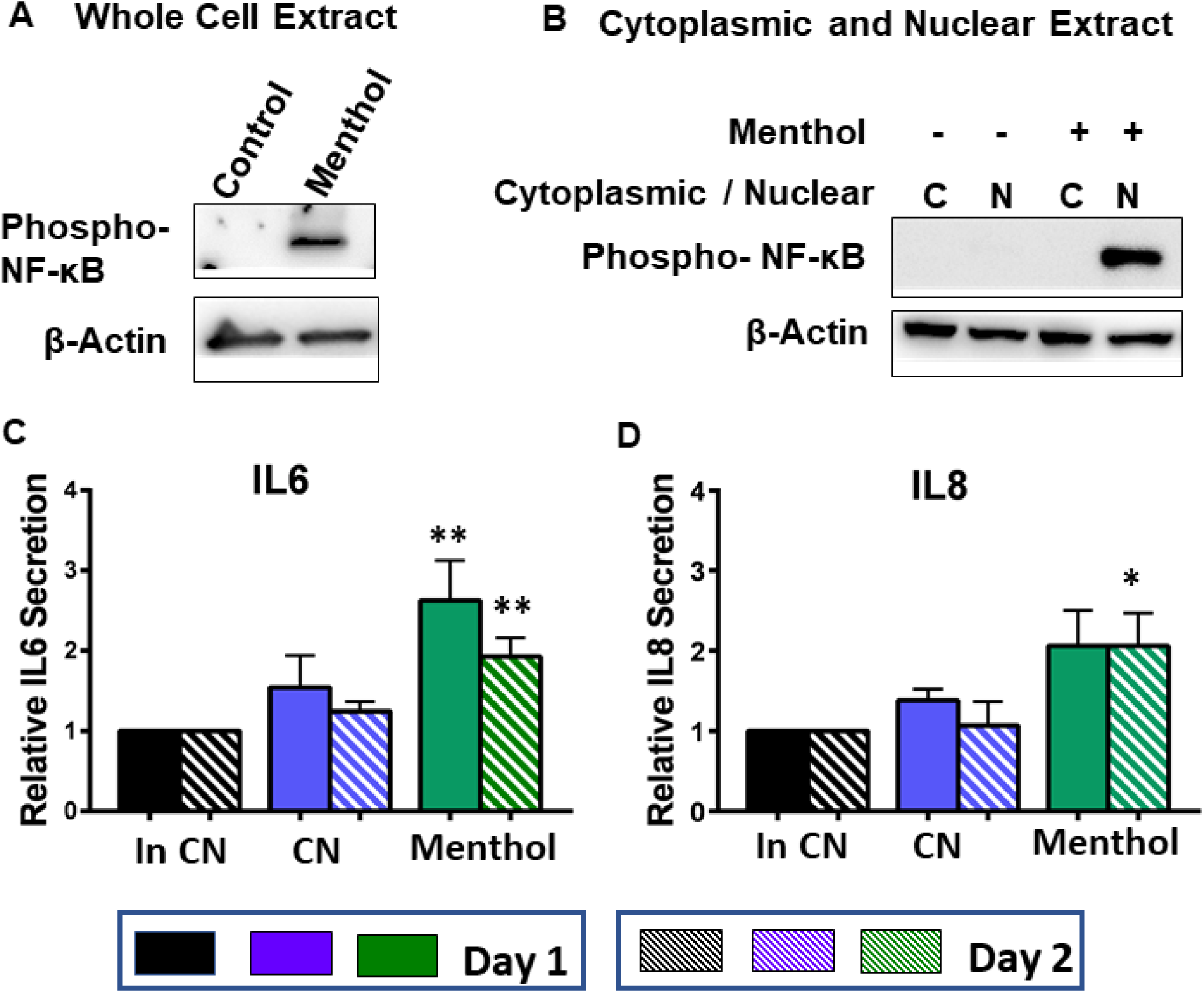
ALI Exposure to Menthol Aerosol in the Cloud Chamber Stimulated Activation of NF-κB and Increased Secretion of Immunomodulatory Cytokines in BEAS-2B cells. **(A and B)** Western blot showing expression of phospho-NF-κB in whole cell extract (A) and in nuclear and cytoplasmic extracts (B) of BEAS-2B cells exposed to menthol aerosol (0.8 mg/mL) in a cloud chamber. β-actin was used as the loading control. **(C and D)** IL-6 levels (Day1 and Day2) and IL-8 (Day1 and Day2) levels in the culture medium, measured by ELISA. Following menthol exposure in the ALI chamber, medium was collected after 24 h (Day 1), replaced with fresh medium, and collected again 24 h later (Day 2). Bars in C and D are the means ± SEM of three independent experiments. Statistical significance was determined using one-way ANOVA with Dunnett’s posthoc test. * = p < 0.05; ** = p < 0.01.

Secretion of Inflammatory Cytokines (IL6 and IL8) is Stimulated by ALI Exposure to Unheated Menthol Aerosol Generated Using a Cloud Chamber IL-6 and IL-8 are cytokines that are up-regulated in inflamed airways and airways of asthma patients (Rincon and Irvin, 2012). The effect of menthol aerosol on secretion of IL6 and IL8 was evaluated following ALI exposure of monolayers of BEAS-2B cells to menthol aerosol in a VITROCELL® cloud chamber. 24 h after exposure, conditioned medium was collected from the inserts, and Day 1 cytokine secretion was analyzed using an ELISA. Fresh medium was added to each insert, and this was collected and analyzed after an additional 24 h of incubation (Day 2). ALI exposure of BEAS-2B cells to unheated menthol aerosol caused an elevation of IL6 and IL8 secretion (Figures 5C, D). Menthol increased the secretion of IL6 and IL8 at least two-fold compared to the control after 24 and 48 h of incubation period.

### Cytotoxic, TEER, and Proteomic Effects of ALI Exposure of EpiAirway Tissue to Heated Menthol Aerosol Produced in an EC

Experiments were next performed using 3D EpiAirway tissues to determine if similar effects on oxidative stress and inflammatory cytokine elevation occurred following ALI exposure to menthol-containing aerosols created using an EC. In this protocol, the aerosol was heated and therefore contained, in addition to menthol, solvent (PG) and any reaction products generated by heating. Endpoints in this protocol included cytotoxicity (MTT assay), TEER measurements, ELISAs, and proteomics analysis of cells following exposure.

EpiAirway tissues were exposed at the ALI to 30 puffs of aerosol produced with an EC at relatively low voltage/power (3V/ 5 watts) then allowed to recover for 24 hours before evaluation with the TEER (Supplementary Figure 2A) and cytotoxicity assays (MTT, LDH) (Supplementary Figure 2B, C). Apart from a small decrease in TEER in the PG control group, tissue integrity was not affected by menthol aerosol treatment when compared to the clean air control (Supplementary Figure 2A). There was no significant effect on mitochondrial reductase activity (Supplementary Figure 2B) in the treatment or PG group. In the LDH assay, there was no effect in the PG control group, and small decrease in the menthol group (Supplementary Figure 2C), which, although statistically significant, may not be biologically relevant.

To determine the effect of menthol aerosol exposures on the proteome of EpiAirway tissue, protein samples were harvested 24 hours after exposure to clean air (CA), PG vehicle control, or menthol. A mass spectrometry (MS) bottom up proteomics method with the False Discovery Rate (FDR) controlled at 1% was performed, which identified 4,462 unique proteins in menthol treated cells (Figures 6A). An in-house statistical method identified 192 significant proteins (35 downregulated and 157 upregulated) in the menthol group and the 22 significant proteins (11 downregulated and 11 upregulated) in the PG group that had differential abundance relative to clean air (Figures 6B, D). Our stringent statistical model was developed (Statistics Analysis Supplemental) to isolate the effect of menthol aerosols from the PG vehicle, resulting in the unconventional shape of the volcano plots (Figures 6B, C). Despite the efforts to exclude PG from the analysis, PG still showed an effect on protein expression (Figure 6C, B), which is consistent with recent reports of PG toxicity and respiratory irritation (Behar et al., 2017; Ghosh et al., 2018)

**Figure 6.**
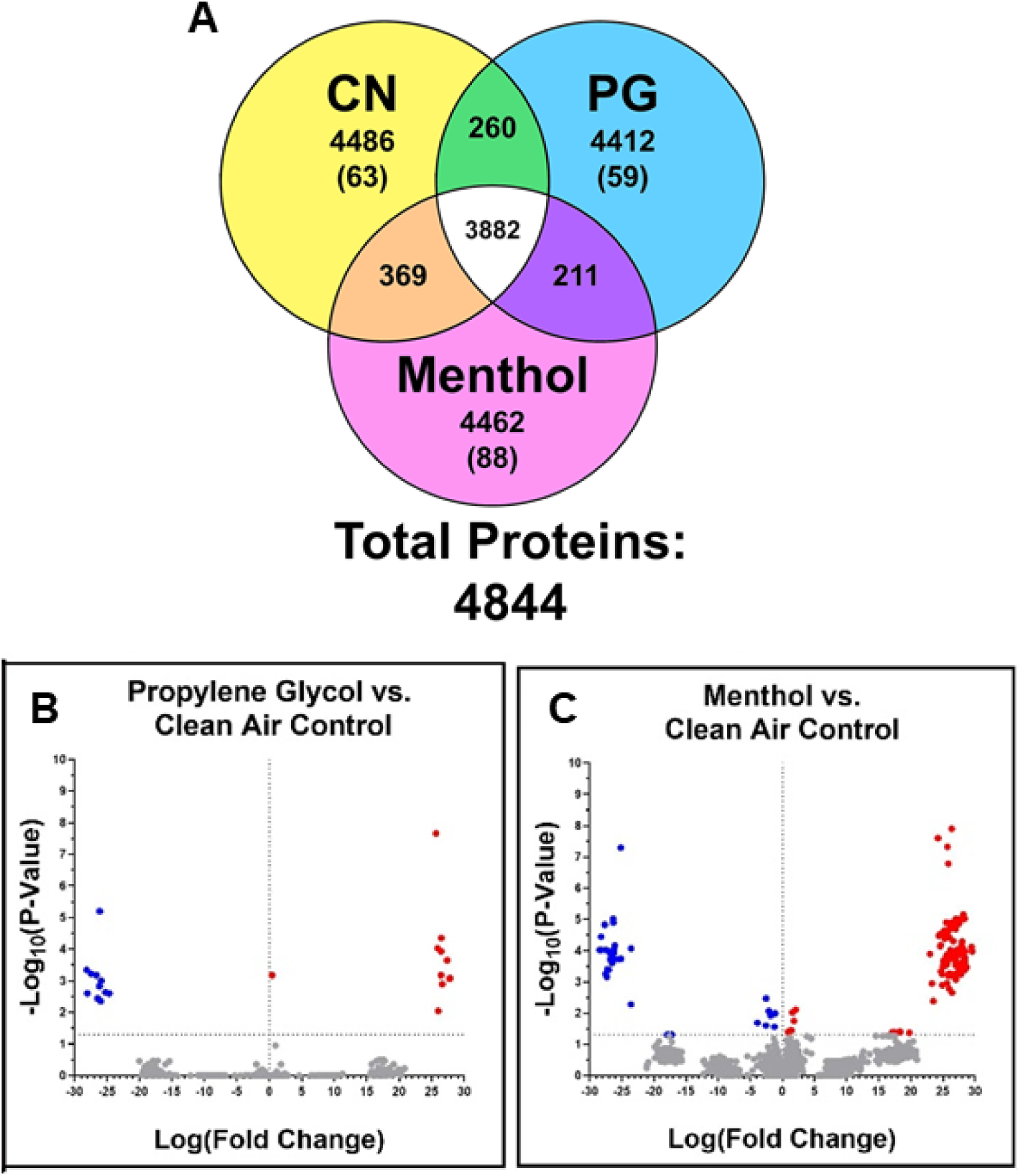
Identification of Proteins Affected by ALI Exposure of EpiAirway Tissue to Menthol or PG Aerosol in the Cultex System. **(A)** Venn diagram of overlapping proteins identified in each treatment group. Values above the parentheses indicate all proteins detected after treatment, while values in parentheses are proteins unique to the CN, PG, and menthol groups. **(B)** Volcano plot showing proteins significantly changed in the PG group relative to the clean air controls. **(C)** Volcano plot showing proteins significantly changed in the menthol group relative to the clean air controls. In B and C, horizontal dashed lines indicate p <0.05. Blue and red dots show down and up regulated proteins, respectively.

### Protein Pathway Interactome Analysis using DAVID

Menthol and PG aerosol exposure data were analyzed using DAVID to show the association of pathway clusters. The color-coding in Figure 7 shows pathway clusters affected by each treatment group (Purple circle: PG, Green circle: Menthol). Menthol aerosol treated cells expressed proteins related to xenobiotic stress, oxidative stress, and inflammation among others, including cytoskeletal activity. Mitochondrial pathway clusters were seen to be affected both by menthol and PG aerosols.

**Figure 7.**
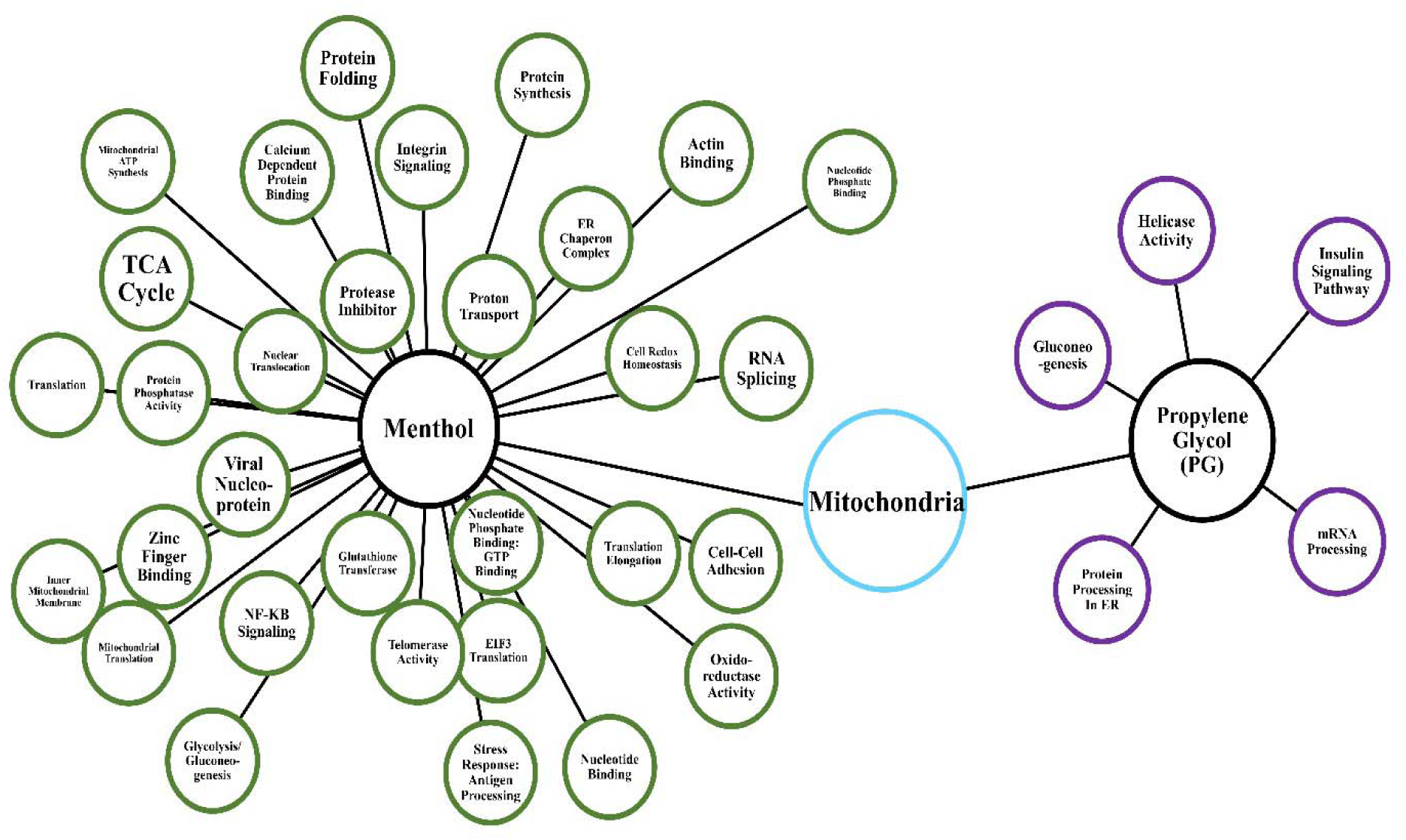
DAVID Derived Interactome of Enrichment Clusters in EpiAirway Tissues Exposed to Menthol or PG at the ALI in the Cultex System. Interaction diagram of proteomics data analyzed with DAVID annotation clustering (p-value <0.05). Only significant proteins with adjusted p-value <0.05 after statistically isolating the effect of PG vehicle were considered for menthol. All proteins significant relative to the Clean Air control (CA) were considered for PG vehicle control.

### Cell Signaling Pathways Affected by Menthol Aerosol Exposure using IPA

IPA pathway enrichment analysis was used to identify canonical pathways significantly impacted by menthol aerosol exposure (Figure 8A). A positive z-score (>2) represents an increase in a cellular process, while a negative z-score (<-2) indicates a decrease. Enrichment of proteins related to oxidative stress (NRF2 mediated oxidative stress response, EIF2 signaling), inflammatory cytokine signaling (IL8 signaling), metabolic pathways (oxidative phosphorylation and gluconeogenesis) among other pathways were found. Top pathways included oxidative phosphorylation (which could increase oxidative stress) and NRF-2 mediated oxidative stress response. Upregulation of EIF2 signaling was verified using western blotting (Supplemental Figure 3A, B). In addition, pathways related to cell proliferation regulation (HIPPO signaling, PTEN signaling, and Cyclins and Cell Cycle Regulation) were downregulated. Chemokine secretion of IL6 and IL8 was investigated using ELISAs and found to be increased significantly in treatment groups relative to clean air controls (Supplemental Figure 4A, B).

**Figure 8.**
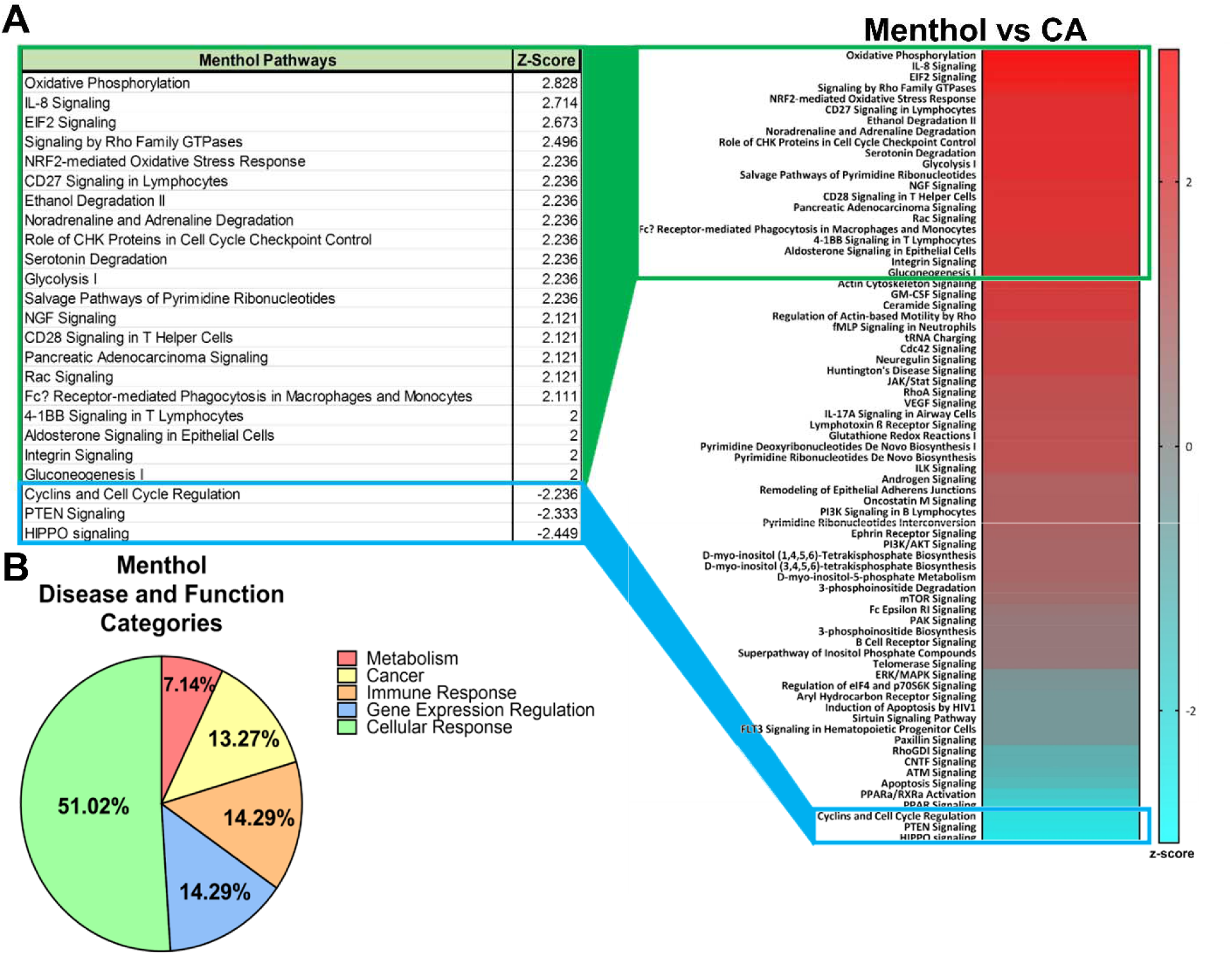
IPA Pathway Analysis and Protein Association Annotations Following 3D Cultex Exposure of EpiAirway at the ALI. **(A)** Heat map of canonical pathways identified with IPA with table of significantly affected pathways (Z-Score>=2; Z-Score<=-2) after Cultex exposure to aerosol from a menthol EC normalized to the Clean Air control (CA). **(B)** Frequency of proteins associated with disease or function identified by IPA in the menthol-treated group. Only proteins with adjusted p-values <0.05 after statistical modeling to isolate the effect of PG vehicle were considered.

Proteins uploaded into IPA (Ingenuity Pathway Analysis) for the menthol group were annotated with associations to various cellular processes. 51.02% of the proteins (N=50) were affiliated with general cellular response, 29% (N=14) with gene expression regulation, 29% (N=14) with immune response, 27% (N=13) with cancer, and 7.14% (N=7) with metabolism pathways (Figure 8B).

## DISCUSSION

To the best of our knowledge, this is the first study to compare the toxic effects of MTT NOAEL concentrations of menthol on human respiratory epithelium using submerged cultures and ALI exposures with and without solvents and with and without heating the aerosols. In most assays, there was excellent agreement of results across the three *in vitro* platforms. At menthol concentrations that did not produce an effect in the MTT assay, oxidative stress was observed with all three platforms, and cytokine elevation/secretion was found in both ALI exposure protocols. These data show that screening toxicants using BEAS-2B cells in 2D submerged cultures or in cloud chamber ALI exposures provides reliable data that could subsequently be confirmed and refined in the more expensive and labor-intensive 3D ALI EpiAirway model. Our data also support the use of submerged cultures for assays that are difficult to perform in 3D ALI exposures, such as monitoring calcium influx through the TRPM8 receptor and live cell imaging.

Menthol induced cytotoxicity in BEAS-2B cells was concentration-dependent (submerged culture protocol). Cytotoxicity (MTT assay) was not observed in the VITROCELL® and Cultex system, probably because exposures were relatively short compared to 24 hours of continual exposure in submerged cultures. In the live cell imaging experiment (submerged culture), the ability of attaching cells to better withstand menthol treatment may be due to removal of cell surface proteins (including TRPM8) by trypsin during detachment of cells for plating (Zhang et al., 2012). Attached cells likely regenerated TRPM8 during the 24-h attachment period before treatment and thus were immediately affected when exposed to menthol. The increased toxicity observed in menthol aerosol fluids during live cell imaging could be due to reaction products, such as formaldehyde, acrolein, and acetaldehyde (Kosmider et al., 2014), that formed from menthol and/or propylene glycol during heating (Behar et al., 2018). In addition, variations in proliferation of attaching vs attached cells in submerged culture show that certain cellular responses can vary within exposure protocols and that cell proliferation is more sensitive to protocol variation than oxidative stress and inflammation. The 3D EpiAirway (Cultex) data on cell proliferation were inconclusive. Some pathways (downregulation of PTEN signaling, downregulation of HIPPO signaling) suggest an increase in cell proliferation, while others (upregulation of CHK, downregulation of cyclins and cell cycle regulation) suggest a decrease (Halder and Johnson, 2011; Jiang and Liu, 2009; Harvey et al., 2013; Stacey, 2003; Wu et al., 2003; Xiao et al., 2006).

We detected the TRPM8 receptor in BEAS-2B cells, A549 cells, and hPFs with relatively more expression in the lung cancer cells (A549). Osteosarcoma, pancreatic, and breast cancer cells also have elevated levels of TRPM8, where it may function in the development and progression of tumors (Liu et al., 2016; Yee, 2015; Zhao et al., 2018). In our immunolabeling data, TRPM8 was localized to the plasma membrane, while another report found it in both the plasma membrane and rough endoplasmic reticulum (ER) (Sabnis et al., 2008). The differences in labeling may be related to the use of different antibodies (polyclonal versus monoclonal). In lung cells, TRPM8 is thought to detect cold temperatures (Bautista et al., 2007), while a less recognized function may be to respond to inhaled chemicals, such as menthol, and activate stress/survival responses.

Our data show that the TRPM8 receptor is functional in BEAS-2B cells. In submerged culture, the initiating event during menthol exposure was a rapid influx of calcium through the TRPM8 receptor, which was inhibited by BCTC. This observation agrees with a previous study in which a higher concentration of menthol (2.5 mM vs 1.3 mM in the current study) induced calcium influx into BEAS-2B cells (Sabnis et al., 2008). Our data showed a rapid increase first in cytosolic calcium (Fig. 1E), suggesting initial influx through the plasma membrane, followed by increased fluorescence in vesicles that are likely of ER origin (Fig.1 F). These vesicles moved adjacent to, but did not fuse with, the plasma membrane, suggesting they quickly sequester excess cytosolic calcium and pump it out near calcium exporters at the cell surface (e.g., Ca^2+^-ATPase and Na+/Ca^2+^ exchanger) (Guerini et al., 2005). It is possible that calcium is pumped out of the ER via a TRPM8 receptor, which has been reported in the ER (Sabnis et al., 2008). Because the TRPM8 receptor would have opposite orientations in the plasma and ER membranes, it is possible TRPM8 in the ER is also activated by menthol and facilitates removal of excess calcium from the cell.

In submerged cultures, menthol also increased mitochondrial ROS in BEAS-2B cells, which was likely due to the increase in intracellular calcium. Elevation of cytosolic calcium can cause a rise in mitochondrial calcium through the mitochondrial uniporter channel (MCU) (Rizzuto et al., 2000; Samanta et al., 2014), and excess calcium in mitochondria can enhance ROS generation (Brookes et al., 2004)

In the VITROCELL® cloud chamber, oxidative stress and an inflammatory response occurred during exposure to relatively low concentrations of menthol (0.8 mg/mL), which did not produce an effect in the cloud chamber MTT assay. The cloud chamber enabled pure menthol-containing aerosol to be tested without solvents (e.g., PG or glycerin) and without heating and reaction-product formation, which distinguishes this protocol from prior ALI studies with EC flavor chemicals (Azzopardi et al., 2016; Leigh et al., 2018). BEAS-2B cells exposed to menthol at the ALI showed an increase in oxidation of mitochondrial proteins and the mitochondrial specific antioxidant enzyme SOD2, both signs of oxidative stress not previously reported for cells treated with menthol at the ALI (Muthumalage et al., 2018; Zhao and Xu, 2016; Zhao et al., 2018). Because these effects were observed in the cloud chamber, they can be attributed to menthol *per se* and not solvents or heat-generated reaction products. Menthol also increased the number of punctate mitochondria, a sign of stress that could lead to mitophagy (Tondera et al., 2009; Zahedi et al., 2018). A similar increase in punctate mitochondria was observed in BEAS-2B and A549 cells treated with rotenone or antimycin, and this change was due to calcium influx into the mitochondria and increased ROS generation (Ahmad et al., 2013). Some of the mitochondrial changes we observed (increased ROS and oxidation of mitochondrial protein) also occurred following exposure of neural stem cells to thirdhand cigarette smoke or electronic cigarette aerosol fluids (Bahl et al., 2016; Zahedi et al., 2018). SOD2, which is located in the mitochondria, is a major ROS detoxifying enzyme (Holley et al., 2011). The elevation of SOD2 in BEAS-2B cells exposed to menthol aerosol was inhibited by siRNA against the TRPM8 receptor, supporting the conclusion that menthol-induced oxidative stress occurred through activation of this receptor.

In the Cultex protocol, 3D EpiAirway tissue was exposed to aerosol which contained pure menthol, PG, and reaction products that formed upon heating the refill fluid in an EC. This aerosol is equivalent to that inhaled by an EC user. Proteins involved in oxidative stress (e.g., oxidative phosphorylation proteins and NRF-2 mediated oxidative stress) and in inflammatory response (e.g., IL-8 signaling) were elevated in the Cultex menthol treated group, consistent with data obtained with the other two exposure protocols. IL-8 signaling, which has been causally linked to acute inflammation (Harada et al., 1994), was the second most upregulated pathway in our Cultex exposure data. While the VITROCELL® cloud chamber data clearly show that menthol by itself can elevate IL8 secretion, the Cultex data further show that PG is also effective, despite efforts to statistically remove its influence from the proteomics analysis. PG is therefore a concern due to its possible adverse health effects (Callahan-lyon, 2014; Wieslander et al., 2001) and ubiquitous use in EC products. EC solvents will be evaluated in more depth in a future study.

In addition to corroborating data obtained with submerged cultures and the cloud chamber, the proteomics analysis of Cultex data identified other pathways that were significantly affected in menthol exposed cells. As examples, NGF signaling, which was increased in the IPA analysis, is involved in activation of NF-κB, a protein that was detected in the DAVID cluster analysis (Figure 7A, 8). NF-κB is normally present in inactive form in cells allowing it to become rapidly activated upon exposure to harmful stimuli (Perkins and Gilmore, 2006). Experiments with tobacco cigarette users and dual users (EC plus and cigarettes) showed upregulation of NGF signaling, glutathione transferase, and NRF2 signaling (D’Anna et al., 2015; Ghosh et al., 2018), suggesting that ECs and conventional cigarettes have similar xenobiotic effects. In addition, Rho Family GTPase signaling, Rac signaling, and integrin signaling are pathways that affect the cytoskeleton (Symons, 1996). Their upregulation may have been involved in the formation of the calcium rich blebs seen in submerged cultures. Blebbing involving these proteins/pathways has been reported in response to calcium influx in human embryonic stem cells upon activation of the P2X7 receptor, which causes rapid influx of calcium (Guan et al., 2016; Weng et al., 2018; Weng and Talbot, 2017).

Our data support the idea that menthol, at concentrations found in EC aerosols, can disturb cell homeostasis and with chronic exposure may contribute to respiratory diseases. Elevation of ROS is involved in numerous diseases, including chronic inflammation (Saito et al., 2006; Takeda et al., 1999; Teramoto et al., 1999). One of the main signaling pathway/transcription factors triggered by oxidative stress is NF-κB (Perkins and Gilmore, 2006). In humans, the bronchiolar epithelium is an important site for NF-κB activation and expression of NF-κB dependent inflammatory mediators (Poynter et al., 2002). NF- κB targets genes that attenuate ROS to promote survival (Djavaheri-Mergny et al., 2004; Kairisalo et al., 2007) and regulates expression of the immunomodulatory cytokines. Elevated NF-κB and induced secretion of two proinflammatory cytokines (IL6 and IL8) are commonly seen in inflammatory pulmonary diseases (Rincon and Irvin, 2012). Acute and chronic inflammation play roles in the pathogenesis of many lung disorders, such as asthma, COPD, adult respiratory distress syndrome, and idiopathic pulmonary fibrosis (Cheng et al., 2007). Although menthol was not established as the causative agent, chronic inflammation from the use of tobacco cigarettes has been linked with Acute Respiratory Distress Syndrome (ARDS) and COPD (Cantin, 2010; Miller et al., 1992; Vaart et al., 2013).

At the high end of the concentration range, menthol is present in some mint flavored EC refill fluids at 84 mg/mL (Behar et al., 2018), which is well above 1 mg/mL, which produces a strong cytotoxic effect in the MTT assay, and the 10 mg/mL that was used for Cultex exposure (Supplemental Fig. 1). Consideration should be given to the possibility that the high concentrations of flavor chemicals in some EC products (Omaiye et al., 2019), such as menthol at 84 mg/mL, could kill the respiratory epithelium resulting in the “burn” characteristics described by some physicians treating EVALI patients (Butt et al., 2019). The dangers of inhaling high concentrations of menthol are further supported by a case report in which acute menthol inhalation caused the death of an other-wise healthy factory worker cleaning a peppermint storage vat; after inhaling menthol fumes for several hours, the 21 year old worker became unconscious, did not respond to treatment, and died 14 days later (Kumar et al., 2016).

At the time of writing, the Center for Disease Control (CDC) reports 2,807 EVALI cases and 68 deaths related to EC usage (Centers for Disease Control and Prevention, 2019). While awaiting firm regulations on the use of flavor chemicals in ECs, the FDA issued a guidance for industry in January 2020 that prohibits the use of flavor chemicals, excluding only tobacco and menthol flavors (Food and Drug Administration Center for Tobacco Products, 2020), potentially leaving public health susceptible to adverse effects from chronic use of menthol at concentrations reported in this study or from acute harm by products with high concentrations of menthol.

## Conclusions

The three *in vitro* platforms for exposing respiratory epithelium to menthol each lead to similar conclusions. Concentrations of menthol within the range found in many EC fluids and aerosols produced rapid calcium influx followed by an increase in oxidative stress and inflammatory cytokines. These responses were inhibited by BCTC and siRNA knock-down of the TRPM8 receptor. Taken together, these data provide a strategy for evaluating the toxicity of inhaled chemicals by first screening in the MTT assay to identify cytotoxic concentrations and possible modes of action. Authentic standards can next be tested at the ALI first using cloud chamber exposure to avoid solvents and reaction products formed by heating, followed by exposure to authentic EC aerosol as done in the Cultex. Using proteomics with ALI exposure systems has the advantage of both confirming and discovering pathways simultaneously. In future studies, it will be valuable to show effects similar to those observed with the EpiAirway protocol in EC users. Validation of the EpiAirway model for translation to *in vivo* exposure would be valuable and could replace animal testing, reduce experimental costs, and accelerate research progress. Data obtained with this approach support the conclusion that menthol, at concentrations found in EC aerosols, adversely affects multiple cell types in the respiratory system, which could disrupt tissue homeostasis, impair cell function, and lead to disease, including some of the recently reported cases of EVALI.

### Limitations of the Study

The effects of menthol were analyzed with exposure to relatively low EC doses. Increasing the number of puffs or voltage of ECs could increase the toxicity of menthol aerosols. In another Cultex study, 200 puffs of aerosol delivered over about 30 min caused differences in cell viability depending on the cell type used (Scheffler et al., 2015). Therefore, results with ALI exposure will vary depending on the protocol. Because of the large variability in EC puffing topography (Behar et al., 2015), it would be useful to develop at least two standard protocols for both the high and low.

## METHODS

All methods can be found in the accompanying “Transparent Methods supplemental file”.

## SUPPLEMENTAL INFORMATION

Supplemental information can be found at …

## Supporting information

Transparent Methods

Statistics Analysis Supplemental

Supplementary Video 1A - Control

Supplementary Video 1B - Menthol

## ACKNOWLEDGEMENTS

Research reported in this publication was supported by NIDA, NIEHS, and the FDA Center for Tobacco Products (CTP) grant #s R01DA036493 and R01ES029741. The content is solely the responsibility of the authors and does not necessarily represent the official views of the NIH or the Food and Drug Administration. The Orbitrap Fusion mass spectrometer was purchased with funds from an NIH shared instrumentation grant (S10OD010669). We thank Lindsey Bustos for her help with the VITROCELL® exposures as well as Man Wong for his helpful remarks.

## AUTHOR CONTRIBUTIONS

Project administration and funding acquisition, P.T and JFP.; Conceptualization, V. N. and P.T.; Investigation, V. N., M T, R.B, Y.W, R.P., W. L and, proteomics analysis, V. N., M. T., S. P., Proteomics statistical analysis, S.Z. and X. C.; Writing, all authors.

---

**Supplementary Figure 1.**
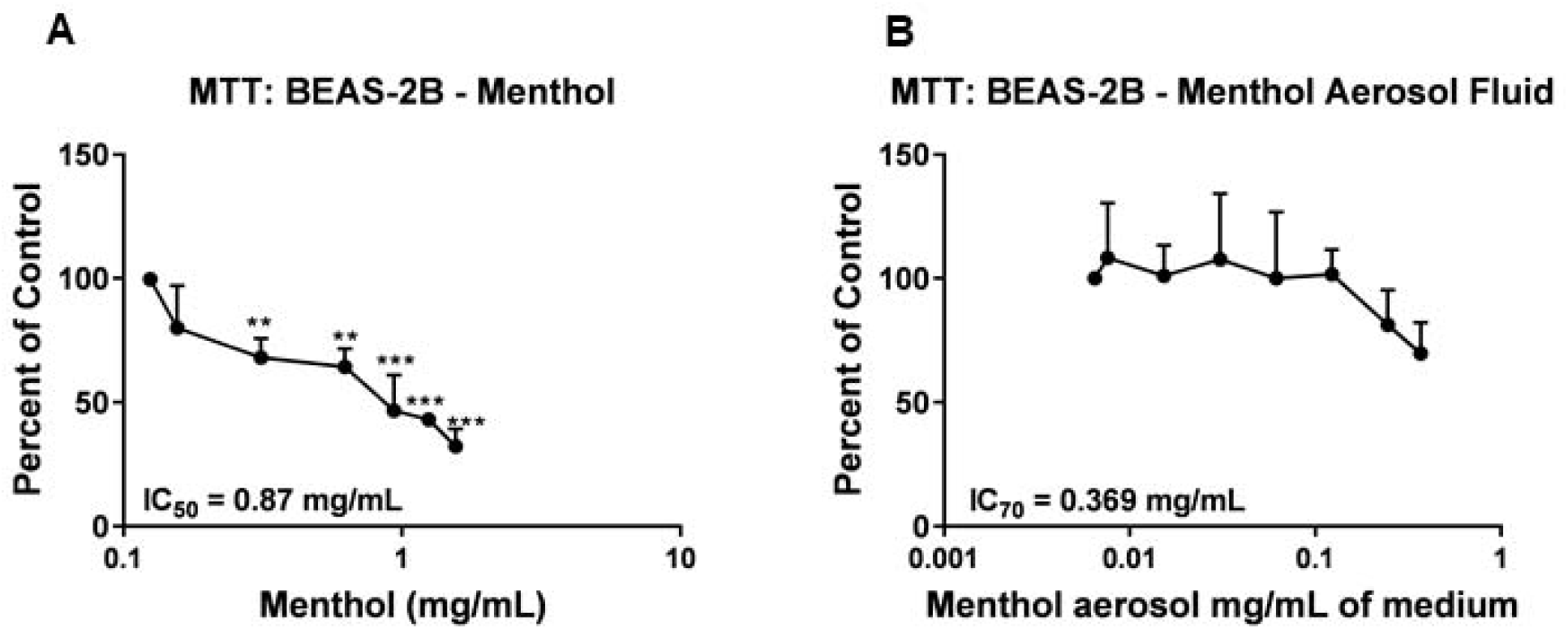
MTT Assay of submerged BEAS-2B cells after exposure to Menthol Treatments. **(A and B)** Analysis of cell metabolism during submerged exposure of BEAS-2B cells using menthol fluid (A) and menthol aerosol fluid (B). Cell metabolism is expressed relative to the control. Data are plotted as means of three independent experiments ± standard error of the mean. Statistical significance was determined with GraphPad Prism using a one-way ANOVA. When significance was found, treated groups were compared with the lowest concentration using Dunnett’s post hoc test. A two-tailed t-test was used to analyze the migration efficiency in the transwell assay. * = p < 0.05; ** = p < 0.01; *** = p<0.001.

**Supplementary Figure 2.**
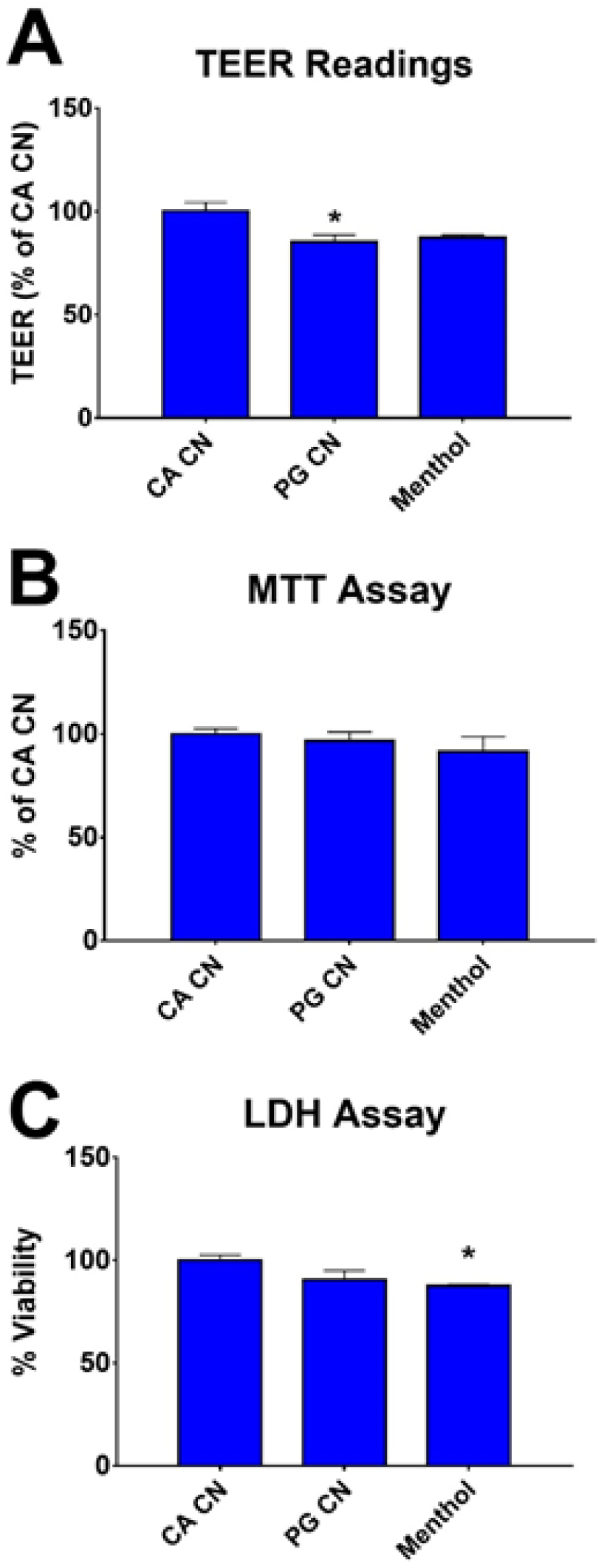
Cytotoxicity Assays on 3D EpiAirway Tissues after exposure to Menthol Aerosols. **(A)** TEER Assay as percent of Clean Air Control. **(B)** MTT Assay as percent of Clean Air Control (C) LDH Assay as percent of Clean Air Control. Statistical significance was determined with GraphPad Prism using a one-way ANOVA for all assays. When significance was found, treated groups were compared with the Clean Air Control using Dunnett’s post hoc test. * = p < 0.05

**Supplementary Figure 3.**
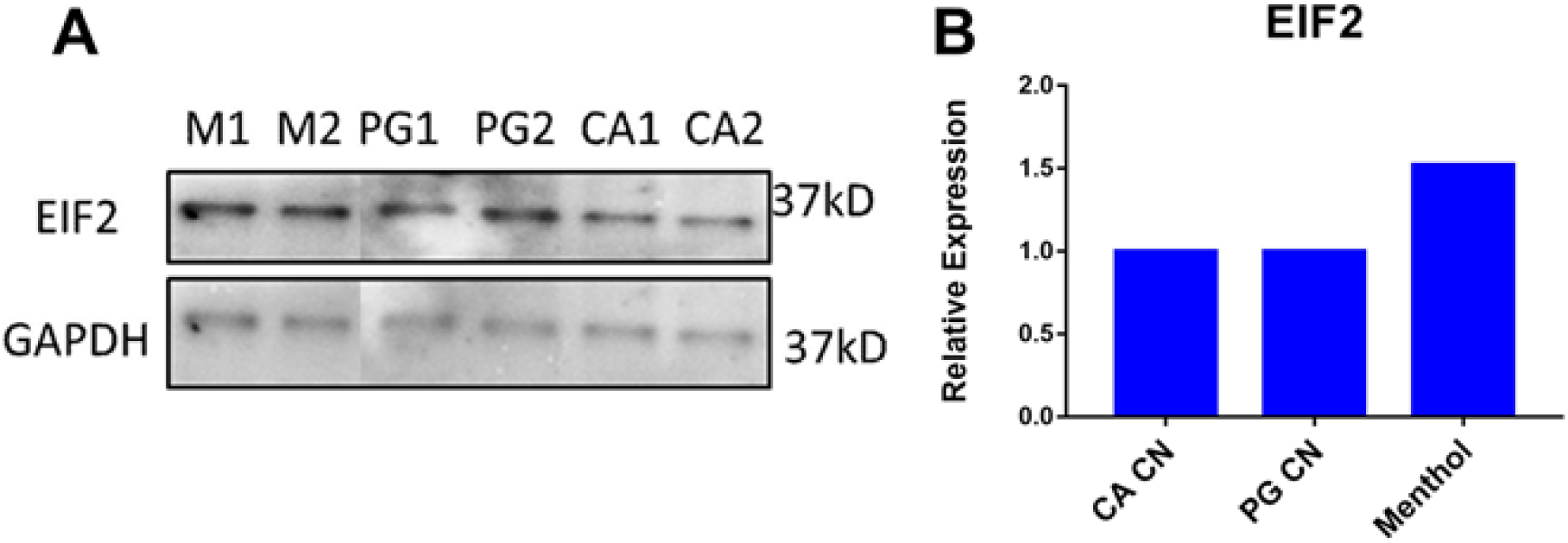
Validation of Proteomic Results on 3D EpiAirway Tissue lysates after exposure in Smoking Machine. **(A)** Western Blot of eIF2 in duplicate after exposure to menthol (M1 and M2), propylene glycol (PG1 and PG 2), or Clean Air (CA1 and CA1) in smoking machine. **(B)** Intensity of western blots quantified with ImageJ and plotted with relative expression to Clean Air control.

**Supplementary Figure 4.**
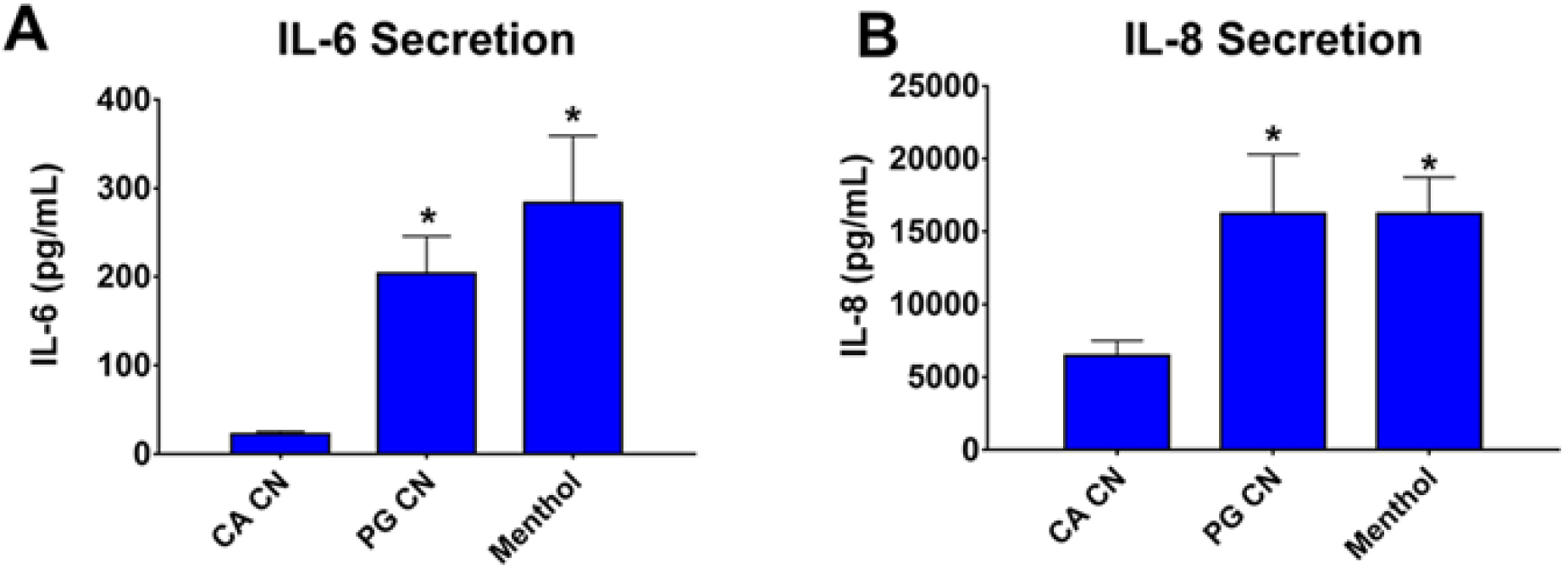
Secretion of cytokines by 3D EpiAirway Tissue into cell media after a 24hr recovery period. **(A)** IL-6 secretion in pg/mL. **(B)** IL-8 Secretion in pg/mL. Statistical significance was determined with GraphPad Prism using a one-way ANOVA for all assays. When significance was found, treated groups were compared with the Clean Air Control using Dunnett’s post hoc test. * = p < 0.05

**Supplementary Video 1. Menthol induced calcium flux: Time-lapse video of BEAS- 2B cells transfected with GCaMP5 showing calcium in control and menthol- treated cells.**

**(A)** 5-minute time course of transfected BEAS-2B cells treated with cell media

**(B)** 5-minute time course of transfected BEAS-2B cells treated with menthol

## References

1. Akerboom, J., Chen, T.-W., Wardill, T.J., Tian, L., Marvin, J.S., Mutlu, S., Calderon, N.C., Esposti, F., Borghuis, B.G., Sun, X.R., et al. (2012). Optimization of a GCaMP calcium indicator for neural activity imaging. J Neurosci. 2012 Oct 3;32(40):13819–40. DOI: 10.1523/JNEUROSCI.2601-12.2012

2. Ahmad, T., Aggarwal, K., Pattnaik, B., Mukherjee, S., Sethi, T., Tiwari, B.K., Kumar, M., Micheal, A., Mabalirajan, U., Ghosh, B., Roy, S.S., Agrawal, A., (2013). Computational classification of mitochondrial shapes reflects stress and redox state. Cell Death Dis. 2013 Jan 17;4: e461. DOI: 10.1038/cddis.2012.213

3. Ai, J., Taylor, K.M., Lisko, J.G., Tran, H., Watson, C.H., Holman, M.R., (2016). Original investigation menthol content in US marketed cigarettes. Nicotine Tob Res. 2016 Jul;18(7):1575–80. https://doi.org/10.1093/ntr/ntv162

4. Azzopardi, D., Patel, K., Jaunky, T., Santopietro, S., Camacho, O.M., Mcaughey, J., Gaça, M., (2016). Electronic cigarette aerosol induces significantly less cytotoxicity than tobacco smoke. Toxicol Mech Methods. 2016 Jul;26(6):477–491. DOI: 10.1080/15376516.2016.1217112

5. Bahl, V., Johnson, K., Phandthong, R., Zahedi, A., Schick, S.F., Talbot, P., (2016). Thirdhand cigarette smoke causes stress-induced mitochondrial hyperfusion and alters the transcriptional profile of stem cells. Toxicol Sci. 2016 Sep;153(1):55–69. DOI: 10.1093/toxsci/kfw102

6. Bahl, V., Lin, S., Xu, N., Davis, B., Wang, Y., Talbot, P., Oak, R., (2012). Comparison of electronic cigarette refill fluid cytotoxicity using embryonic and adult models. Reprod Toxicol. 2012 Dec;34(4):529–37. DOI: 10.1016/j.reprotox.2012.08.001

7. Barrington-Trimis, J.L., Leventhal, A., (2018). Adolescents’ use of “pod mod” e-cigarettes - urgent concerns. N Engl J Med. 2018 Sep 20;379(12):1099–1102. DOI: 10.1056/NEJMp1805758

8. Bautista, D.M., Siemens, J., Glazer, J.M., Tsuruda, P.R., Basbaum, A.I., Stucky, C.L., Jordt, S., Julius, D., (2007). The menthol receptor TRPM8 is the principal detector of environmental cold. Nature. 2007 Jul 12;448(7150):204–8. DOI: 10.1038/nature05910

9. Behar, R.Z., Hua, M., Talbot, P., (2015). Puffing topography and nicotine intake of electronic cigarette users. PLoS One. 2015 Feb 9;10(2):e0117222. DOI:10.1371/journal.pone.0117222

10. Behar, R.Z., Luo, W., Mcwhirter, K.J., Pankow, J.F., Talbot, P., (2018). Analytical and toxicological evaluation of flavor chemicals in electronic cigarette refill fluids. Scientific Reports volume 8, Article number: 8288 (2018). DOI: 10.1038/s41598-018-25575-6

11. Behar, R.Z., Wang, Y., Talbot, P., (2017). Comparing the cytotoxicity of electronic cigarette fluids, aerosols and solvents. Tob Control. 2018 May;27(3):325–333. DOI: 10.1136/tobaccocontrol-2016-053472

12. Brookes, P.S., Yoon, Y., Robotham, J.L., Anders, M.W., Sheu, S., (2004). Calcium, ATP, and ROS: a mitochondrial love-hate triangle. Am J Physiol Cell Physiol. 2004 Oct;287(4):C817–33. DOI: 10.1152/ajpcell.00139.2004

13. Butt, Y.M., Smith, M.L., Tazelaar, H.D., Vaszar, L.T., Swanson, K.L., Cecchini, M.J., Boland, J.M., Bois, M.C., Boyum, J.H., Froemming, A.T., Khoor, A., Mira-Avendano, I., Patel, A., Larsen, B.T., (2019). Pathology of Vaping-Associated Lung Injury. N Engl J Med 2019; 381:1780–1781. DOI: 10.1056/NEJMc1913069

14. Callahan-lyon, P., (2014). Electronic cigarettes: human health effects. Tobacco control 23 Suppl 2(Suppl 2): ii36–ii40· May 2014. http://dx.doi.org/10.1136/tobaccocontrol-2013-051470

15. Cantin, A.M., (2010). Cellular response to cigarette smoke and oxidants: adapting to survive. Proc Am Thorac Soc. 2010 Nov;7(6):368–75. DOI: 10.1513/pats.201001-014AW

16. Cheng, D., Han, W., Chen, S.M., Taylor, P., Chont, M., Park, G., Sheller, J.R., Polosukhin, V. V, Christman, J.W., Yull, F.E., Blackwell, T.S., Cheng, D., Han, W., Chen, S.M., Sherrill, T.P., Chont, M., Park, G., Sheller, J.R., Polosukhin, V. V, Christman, J.W., Yull, F.E., Blackwell, T.S., (2007). Through the NF- κB pathway 1. J Immunol May 15, 2007, 178 (10) 6504–6513. DOI: https://doi.org/10.4049/jimmunol.178.10.6504

17. D’Anna, C., Cigna, D., Costanzo, G., Bruno, A., Ferraro, M., Vincenzo, D., Bianchi, L., Bini, L., Pace, E., (2015). Cigarette smoke alters the proteomic profile of lung fibroblasts. Mol Biosyst. 2015 Jun;11(6):1644–52. DOI: 10.1039/c5mb00188a

18. Delnevo, C.D., Gundersen, D.A., Hrywna, M., Echeverria, S.E., Steinberg, M.B., (2011). Smoking-cessation prevalence among U.S. smokers of menthol versus non-menthol cigarettes. Am J Prev Med. 2011 Oct;41(4):357–65. DOI: 10.1016/j.amepre.2011.06.039

19. DeVito, E.E., Jensen, K.P., O’Malley, S.S., Gueorguieva, R., Krishnan-Sarin, S., Valentine, G., Jatlow, P.I., Sofuoglu, M., (2019). Modulation of “protective” nicotine perception and use profile by flavorants: preliminary findings in e-cigarettes. Nicotine Tob Res. 2019 Apr 17. pii: ntz057. DOI: 10.1093/ntr/ntz057

20. Djavaheri-Mergny, M., Javelaud, D., Wietzerbin, J., Besancon, F., (2004). NF-κB activation prevents apoptotic oxidative stress via an increase of both thioredoxin and MnSOD levels in TNF α-treated Ewing sarcoma cells. FEBS Lett. 2004 Nov 11;578(1-2): 111–115. DOI: https://doi.org/10.1016/j.febslet.2004.10.082

21. Ending new nicotine dependencies act of 2019 (ENND Act), S.2519, 116th cong. (2019- 2020).

22. Family Smoking Prevention and Tobacco Control Act of 2009, 5 USC §§ 1776-1858 (2009).

23. Food and Drug Administration, (2011). Preliminary scientific evaluation of the possible public health effects of menthol versus nonmenthol cigarettes. Retrieved from https://permanent.access.gpo.gov/gpo39032/Preliminary%20Scientific%20Evaluation%20Menthol%20508%20reduced.pdf

24. Food and Drug Administration, Center for Tobacco Products, (2020). Enforcement Priorities for Electronic Nicotine Delivery Systems (ENDS) and Other Deemed Products on the Market Without Premarket Authorization. Retrieved from: https://www.fda.gov/regulatory-information/search-fda-guidance-documents/enforcement-priorities-electronic-nicotine-delivery-system-ends-and-other-deemed-products-market

25. Ghosh, A., Coakley, R.C., Mascenik, T., Rowell, T.R., Davis, E.S., Rogers, K., Webster, M.J., Dang, H., Herring, L.E., Sassano, M.F., Livraghi-butrico, A., Buren, S.K. Van, Graves, L.M., Herman, M.A., Randell, S.H., Alexis, N.E., Tarran, R., Carolina, N., (2018). Chronic e-cigarette exposure alters the human bronchial epithelial proteome. Am J Respir Crit Care Med. 2018 Jul 1;198(1):67–76. DOI: 10.1164/rccm.201710-2033OC

26. Grady, D., (2019, Nov 8). Lung damage from vaping resembles chemical burns, report says. The New York Times

27. Guan, B.X., Bhanu, B., Talbot, P., Weng, N.J., (2016). Extraction of blebs in human embryonic stem cell videos. IEEE/ACM Trans Comput Biol Bioinform. 2016 Jul-Aug;13(4):678–88. DOI: 10.1109/TCBB.2015.2480091

28. Guerini, D., Coletto, L., Carafoli, E., (2005) Exporting calcium from cells. Cell Calcium 38: 281–289.

29. Halder, G., Johnson, R.L., (2011). Hippo signaling: growth control and beyond. Development. 2011 Jan;138(1):9–22. DOI: 10.1242/dev.045500

30. Hallagan, J., (2014). The safety assessment and regulatory authority to use flavors: focus on e-cigarettes. Retrieved from: https://www.femaflavor.org/node/24344

31. Harada, A., Sekido, N., Akahoshi, T., Wada, T., Mukaida, N., Matsushima, K., (1994). Essential involvement of interleukin-8 (IL-8) in acute inflammation. J Leukoc Biol. 1994 Nov;56(5):559–64. https://doi.org/10.1002/jlb.56.5.559

32. Harvey, K.F., Zhang, X., Thomas, D.M., (2013). The hippo pathway and human cancer. Nat Rev Cancer. 2013 Apr;13(4):246–57. DOI: 10.1038/nrc3458

33. Henderson, B.J., (2019). Chapter 12 - linking nicotine, menthol, and brain changes. In Neuroscience of Nicotine Mechanisms and Treatment, V. Preedy, ed. (Science Direct), pp. 87–95.

34. Holley, A.K., Bakthavatchalu, V., Velez-roman, J.M., Clair, D.K.S., (2011). Manganese superoxide dismutase: guardian of the powerhouse. Int J Mol Sci. 2011;12(10):7114–62. DOI: 10.3390/ijms12107114

35. Hua, M., Omaiye, E.E., Luo, W., Mcwhirter, K.J., Pankow, J.F., Talbot, P., (2019). Identification of cytotoxic flavor chemicals in top-selling electronic cigarette refill fluids. Scientific Reports volume 9, Article number: 2782 (2019). DOI:10.1038/s41598-019-38978-w

36. Jiang, B., Liu, L., (2009). PI3K / PTEN signaling in tumorigenesis and angiogenesis. Adv Cancer Res. 2009;102:19–65. DOI: 10.1016/S0065-230X(09)02002-8

37. Kairisalo, M., Korhonen, L., Blomgren, K., Lindholm, D., (2007). X-linked inhibitor of apoptosis protein increases mitochondrial antioxidants through NF- κB activation. Biochem Biophys Res Commun. 2007 Dec 7;364(1):138–44. DOI: 10.1016/j.bbrc.2007.09.115

38. Kosmider, L., Sobczak, A., Fik, M., Knysak, J., Zaciera, M., Kurek, J., Goniewicz, M.L., (2014). Carbonyl compounds in electronic cigarette vapors: effects of nicotine solvent and battery output voltage. Nicotine Tob Res. 2014 Oct;16(10):1319–26. DOI: 10.1093/ntr/ntu078

39. Kumar, A., Baitha, U., Aggarwal, P., and Jammshed, N. (2016) A fatal case of menthol poisoning. Int Journal of App and Basic Med Res. 6: 137–139.

40. Laker, R.C., Xu, P., Ryall, K.A., Sujkowski, A., Kenwood, B.M., Chain, K.H., Zhang, M., Royal, M.A., Hoehn, K.L., Driscoll, M., Adler, P.N., Wessells, R.J., Saucerman, J.J., Yan, Z., (2014). A novel mitotimer reporter gene for mitochondrial content, structure, stress, and damage in vivo. J Biol Chem. 2014 Apr 25;289(17):12005–15. DOI: 10.1074/jbc.M113.530527

41. Leigh, N.J., Lawton, R.I., Hershberger, P.A., Goniewicz, M.L., (2018). Flavorings significantly affect inhalation toxicity of aerosol generated from electronic nicotine delivery systems (ENDS). Tobacco Control, 25(Suppl 2), ii81, November 2016. DOI:10.1136/tobaccocontrol-2016-053205

42. Lin, A.-H., Liu, M., Ko, H., Perng, D., Lee, T., Kou, Y.R., (2018). Menthol cigarette smoke induces more severe lung inflammation than non-menthol cigarette smoke does in mice with subchronic exposure – role of TRPM8. Front Physiol. 2018 Dec 18;9:1817. DOI: 10.3389/fphys.2018.01817

43. Lisko, J., Stanfill, S., Watson, C., (2014). Analytical methods tobacco products. Anal. Methods 6, 4698–4704.

44. Liu, Z., Wu, H., Wei, Z., Wang, X., Shen, P., (2016). TRPM8: a potential target for cancer treatment. J Cancer Res Clin Oncol. 2016 Sep;142(9L):1871–81. DOI: 10.1007/s00432-015-2112-1

45. Lloyd, J., O’Malley, P., Miech, R., Bachman, J., Schulenberg, J., (2019). Monitoring the Future National Survey Results on Drug Use, 1975-2015: Overview, Key Findings on Adolescent Drug Use. Retrieved from https://eric.ed.gov/?id=ED578539

46. Miller, E.J., Cohen, A.B., Nagao, S., Griffith, D.E., Maunder, R.J., Martin, T.R., Wiener-Kronish, J.P., Sticherling, M., Chrisophers, E., Matthay, M.A., (1992). Elevated levels of NAP-1/Interleukin-8 are present in the airspaces of patients with the adult respiratory distress syndrome and are associated with increased mortality. Am Rev Respir Dis. 1992 Aug;146(2):427–32. DOI: 10.1164/ajrccm/146.2.427

47. Muthumalage, T., Prinz, M., Ansah, K.O., Gerloff, J., Sundar, I.K., Rahman, I., (2018). Inflammatory and oxidative responses induced by exposure to commonly used e-cigarette flavoring chemicals and flavored e-liquids without nicotine. Front Physiol. 2018 Jan 11;8:1130. DOI: 10.3389/fphys.2017.01130

48. Center for Disease Control and Prevention, (2019). States update number of hospitalized EVALI cases and EVALI deaths. Retrieved from https://www.cdc.gov/media/releases/2019/s1231-evali-cases-update.html

49. Nonnemaker, J., Hersey, J., Homsi, G., Busey, A., Allen, J., Vallone, D., (2013). Initiation with menthol cigarettes and youth smoking uptake. Addiction. 2013 Jan;108(1):171–8. DOI: 10.1111/j.1360-0443.2012.04045.x

50. Omaiye, E.E., Mcwhirter, K.J., Luo, W., Peyton A., Pankow, J.F., Talbot, P. (2019) High concentrations of flavor chemicals are present in electronic cigarette refill fluids. Scientific Reports 9: 2468. https://doi.org/10.1038/s41598-019-39550-2

51. Omaiye, E.E., Mcwhirter, K.J., Luo, W., Pankow, J.F., Talbot, P., (2018). Toxicity of JUUL fluids and aerosols correlates strongly with nicotine and some flavor chemical concentrations. Chem Res Toxicol. 2019 Jun 17;32(6):1058–1069. DOI: 10.1021/acs.chemrestox.8b00381

52. Paschke, M., Tkachenko, A., Ackermann, K., Hutzler, C., Henkler, F., Luch, A., (2017). Activation of the cold-receptor TRPM8 by low levels of menthol in tobacco products. Toxicol Lett. 2017 Apr 5;271:50–57. DOI: 10.1016/j.toxlet.2017.02.020

53. Peier, A.M., Moqrich, A., Hergarden, A.C., Reeve, A.J., Andersson, D.A., Story, G.M., Earley, T.J., Dragoni, I., Mcintyre, P., Bevan, S., Patapoutian, A., Diego, S., (2002). A TRP channel that senses cold stimuli and menthol. Cell. 2002 Mar 8;108(5):705–15. DOI:10.1016/s0092-8674(02)00652-9

54. Perkins, N.D., Gilmore, T.D., (2006). Good cop, bad cop: the different faces of NF- KB. Cell Death Differ. 2006 May;13(5):759–72. DOI: 10.1038/sj.cdd.4401838

55. Poynter, M.E., Irvin, C.G., Janssen-Heininger, Y.M.W., (2002). Rapid activation of nuclear factor- κB in airway epithelium in a murine model of allergic airway inflammation. Am J Pathol. 2002 Apr;160(4):1325–34. DOI: 10.1016/s0002-9440(10)62559-x

56. Rincon, M., Irvin, C.G., (2012). Role of IL-6 in asthma and other inflammatory pulmonary diseases. Int J Biol Sci. 2012; 8(9): 1281–1290. doi: 10.7150/ijbs.4874

57. Rizzuto, R., Bernardi, P., Pozzan, T., (2000). Topical review mitochondria as all-round players of the calcium game. J Physiol. 2000 Nov 15;529 Pt 1:37–47. DOI: 10.1111/j.1469-7793.2000.00037.x

58. Sabnis, A.S., Shadid, M., Yost, G.S., Reilly, C.A., (2008). Human lung epithelial cells express a functional cold-sensing TRPM8 variant. Am J Respir Cell Mol Biol. 2008 Oct;39(4):466–74. DOI: 10.1165/rcmb.2007-0440OC

59. Saito, Y., Nishio, K., Ogawa, Y., Kimata, J., Kinumi, T., Yoshida, Y., Noguchi, N., Niki, E., (2006). Turning point in apoptosis/necrosis induced by hydrogen peroxide. Free Radic Res. 2006 Jun;40(6):619–30. DOI: 10.1080/10715760600632552

60. Samanta, K., Douglas, S., Parekh, A.B., (2014). Mitochondrial calcium uniporter MCU supports cytoplasmic Ca2+ oscillations, store-operated Ca2+ entry and Ca2+ - dependent gene expression in response to receptor stimulation. PLoS ONE (Vol. 9, Issue 7) https://doi.org/10.1371/journal.pone.0101188

61. Scheffler, S., Dieken, H., Krischenowski, O., Aufderheide, M., (2015). Cytotoxic evaluation of e-liquid aerosol using different lung-derived cell models. Int J Environ Res Public Health. 2015 Oct; 12(10): 12466–12474. DOI: 10.3390/ijerph121012466

62. Stacey, D.W., (2003). Cyclin D1 serves as a cell cycle regulatory switch in actively proliferating cells. Curr Opin Cell Biol. 2003 Apr;15(2):158–63. DOI: 10.1016/s0955-0674(03)00008-5

63. Symons, M., (1996). Rho family GTPases Biochem Sci. 1996 May;21(5):178–81. DOI: https://doi.org/10.1016/S0968-0004(96)10022-0

64. Takeda, K., Clausen, B.E., Kaisho, T., Tsujimura, T., Terada, N., Forster, I., Akira, S., (1999). Enhanced Th1 activity and development of chronic enterocolitis in mice devoid of Stat3 in macrophages and neutrophils. Immunity. 1999 Jan;10(1):39–49. DOI: 10.1016/s1074-7613(00)80005-9

65. Teramoto, S., Tomita, T., Matsui, H., Ohga, E., Matsuse, T., Ouchi, Y., (1999). Hydrogen peroxide-induced apoptosis and necrosis in human lung fibroblasts: protective roles of glutathione. Jpn J Pharmacol. 1999 Jan;79(1):33–40. DOI: 10.1254/jjp.79.33

66. Tierney, P.A., Karpinski, C.D., Brown, J.E., Luo, W., Pankow, J.F., (2016). Flavour chemicals in electronic cigarette fluids. Tob Control. 2016 Apr;25(e1):e10-5. DOI: 10.1136/tobaccocontrol-2014-052175

67. Tondera, D., Jourdain, A., Karbowski, M., Mattenberger, Y., Cruz, S. Da, Clerc, P., Raschke, I., Merkwirth, C., Ehses, S., Krause, F., Chan, D.C., Alexander, C., Bauer, C., Youle, R., Langer, T., Martinou, J., (2009). SLP-2 is required for stress-induced mitochondrial hyperfusion. EMBO J. 2009 Jun 3;28(11):1589–600. DOI: 10.1038/emboj.2009.89

68. U.S. Department of Health and Human Services. (2016). E-Cigarette Use Among Youth and Young Adults. A Report of the Surgeon General. Atlanta, GA: U.S. Department of Health and Human Services, Centers for Disease Control and Prevention, National Center for Chronic Disease Prevention and Health Promotion, Office on Smoking and Health, 2016.

69. Vaart, H.J.D., Murgatroyd, S., Rossiter, H.B., Chen, C., Casaburi, R., Porszasz, J., (2013). Using the power-duration curve to select constant work rates for endurance testing in COPD. American Journal of Respiratory and Critical Care Medicine 2013; 187: A1365

70. Villanti, A.C., Collins, L.K., Niaura, R.S., Gagosian, S.Y., Abrams, D.B., (2017). Menthol cigarettes and the public health standard: a systematic review. BMC Public Health. 2017 Dec 29;17(1):983. DOI: 10.1186/s12889-017-4987-z

71. Weng, N.J., Cheung, C., Talbot, P., (2018). Dynamic blebbing: A bottleneck to human embryonic stem cell culture that can be overcome by Laminin-Integrin signaling. Stem Cell Res. 2018 Dec; 33:233–246. DOI: 10.1016/j.scr.2018.10.022

72. Weng, N.J., Talbot, P., (2017). The P2X7 receptor is an upstream regulator of dynamic blebbing and a pluripotency marker in human embryonic stem cells. Stem Cell Res. 2017 Aug; 23:39-49. DOI: 10.1016/j.scr.2017.06.007

73. Wieslander, G., Norbäck, D., Lindgren, T., (2001). Experimental exposure to propylene glycol mist in aviation emergency training: acute ocular and respiratory effects. Occup Environ Med. 2001 Oct; 58(10): 649–655. DOI: 10.1136/oem.58.10.649

74. Willis, D.N., Liu, B., Ha, M.A., Jordt, S., Morris, J.B., (2011). Menthol attenuates respiratory irritation responses to multiple cigarette smoke irritants. FASEB J. 2011 Dec;25(12):4434–44. DOI: 10.1096/fj.11-188383

75. Wu, H., Goel, V., Haluska, F.G., (2003). PTEN signaling pathways in melanoma. Oncogene. 2003 May 19;22(20):3113–22. DOI: 10.1038/sj.onc.1206451

76. Xiao, Z., Xue, J., Sowin, T.J., Zhang, H., (2006). Differential roles of checkpoint kinase 1, checkpoint kinase 2, and mitogen-activated protein kinase-activated protein kinase 2 in mediating DNA damage-induced cell cycle arrest: Implications for cancer therapy. Molecular Cancer Therapeutics 5(8):1935–43. DOI: 10.1158/1535-7163.MCT-06-0077.

77. Yee, N.S., (2015). Roles of TRPM8 ion channels in cancer: proliferation, survival, and invasion. Cancers (Basel). 2015 Oct 23; 7 (4):2134–46. DOI: 10.3390/cancers7040882

78. Zahedi, A., Phandthong, R., Chaili, A., Remark, G., Talbot, P., (2018). Epithelial-to-mesenchymal transition of A549 lung cancer cells exposed to electronic cigarettes. Lung Cancer. 2018 Aug;122:224–233. DOI: 10.1016/j.lungcan.2018.06.010

79. Zhang, B., Shan, H., Li, D., Li, Z., Zhu, K., (2012). Different methods of detaching adherent cells significantly affect the detection of TRAIL receptors. Tumori 98, 800–803. DOI: 10.1700/1217.13506

80. Zhao, W., Xu, H., (2016). High expression of TRPM8 predicts poor prognosis in patients with osteosarcoma. Oncol Lett. 2016 Aug;12(2):1373–1379. DOI: 10.3892/ol.2016.4764

81. Zhao, Y., Du, L., Du, G., (2018). Menthol. natural small molecule drugs from plants [981-10-8021-6; 981-10-8022-4] pg:289–294

