## Supplementary material for "Menthol in Electronic Cigarettes: A Contributor to Respiratory Disease?": Transparent Methods

**MATERIALS and METHODS**

**Chemicals**

Menthol, BCTC (N-(4-tert.-butyl-phenyl)-4-(3-chloropyridin-2-yl) tetrahydropyrazine-1(2H)-carboxamide) and siRNA oligonucleotide against TRPM8 were purchased from Sigma (St. Louis, MO). Bronchial epithelial growth medium (BEGM) was purchased from Lonza (Walkersville, MD). TRPM8 antibody was purchased from Abcam (Cambridge, MA). SOD2 and β-actin antibody were purchased from Cell Signaling (Danvers, MA). MitoSOX dye, Nuclear and Cytoplasmic extraction kit (Catalog #78833), and Lipofectamine RNAiMAX Transfection Reagent were purchased from Thermo Fisher Scientific (Waltham, MA). Mitotimer and GCaMP5 plasmids were purchased from Addgene (Cambridge, MA).

**Cell culture and exposure:**

This study utilized both submerged culture and ALI exposure. Cell viability, proliferation, live cell imaging of calcium influx, and immunodetection of the TRPM8 receptor were carried out in submerged cultures using BEAS-2B cells. Cell viability, inflammatory response, and oxidative stress were carried out using ALI exposure in either a cloud chamber (BEAS-2B cells) or Cultex exposure system (EpiAirway).

For submerged cultures, human bronchial epithelial cells (BEAS-2B) (American Type Culture Collection, Rockville, MD) were expanded and grown in serum free BEGM supplemented with growth factors. Culture ﬂasks (Corning, Inc, Corning, NY) were pre-coated with BEBM (Bronchial Epithelial Basal Medium) fortiﬁed with collagen (30 mg/ml), ﬁbronectin (10 mg/ml), and bovine serum albumin (BSA, 10 mg/ml). Cells were maintained at 37^o^ C, between 30 and 90% conﬂuence, in a humidiﬁed incubator with 5% carbon dioxide. For subculturing, cells were trypsin-dissociated and passaged every 2 to 4 days.

For ALI exposure in the VITROCELL® cloud chamber, 12 mm transwell inserts with a pore size of 0.4µm (Corning NY, USA) were pre-coated with BEBM medium. BEAS-2B cells were plated at 60,000 cells per insert in BEGM medium. After 48 h, basal medium was replaced with fresh medium, and medium from the apical layer was removed to enable ALI culture. The apical layer of the transwells was washed twice with 0.5mL phosphate-buffered saline (PBS) immediately before exposure to aerosol in the cloud chamber.

ALI experiments in the Cultex exposure system were done using 3D human EpiAirway tissues purchased from Mat-Tek Corp (Ashland, MA). EpiAirway is a 3D mucociliary tissue model consisting of normal, human bronchial epithelial cells (hBEC). EpiAirway was prepared from primary hBECs isolated at Mat-Tek Corp from a 23-year-old Caucasian male non-smoker with no history of respiratory disease. Tissues were obtained by Mat-Tek for research purposes with informed consent. After receiving EpiAirway inserts in 12-well plates, the tissues were equilibrated in 700 µL of AIR-100-ASY assay medium/well for at least 24 hours at 37^o^C with 5% CO_2_ before ALI exposures.

**Aerosol generation for exposure of submerged cultures**

For submerged culture exposures, aerosol fluids were made in BEAS-2B culture medium as described previously (Behar et al., 2016, 2014). Menthol aerosols were produced with fresh unused Vea cartomizers at 2.9 V, 2.1 Ω, and 4W. Transference to aerosols, which was not measured, was assumed to be 100%. Lab made refill fluid (6 mL) containing 10 mg/mL of menthol was prepared in 80% propylene glycol and 20% distilled water, and 1mL was loaded into each cartomizer as recommended by the vendor. Puff duration was 4.3 s (Behar et al., 2015), and flow rate was adjusted to produce consistent robust clouds of aerosol which were collected in a round-bottom flask submerged in an ice bath containing BEAS-2B culture medium. Aerosol solutions were made at a concentration of six total puff equivalents (TPE), where one TPE = the number of puffs fully dissolved in 1 mL of culture medium. For each batch, 18 puffs were collected in 3 mL of medium placed in a round bottom flask in an ice bath.

**Menthol aerosol exposure using the VITROCELL® Cloud chamber**

BEAS-2B cells cultured in transwell inserts were placed in a VITROCELL® Cloud exposure system (Waldkirch, Germany) designed for spatially uniform deposition of aerosols. Stock solutions containing various amounts of menthol (0.468 g/mL, 0.156 g/mL, and 0.078 g/mL) were made in dimethyl sulfoxide (DMSO). Working solutions of menthol ranging from 0.15 mg/mL to 3 mg/mL were made immediately before use by dissolving the stock solutions in 0.8% NaCl. 200 µL of the working solutions with or without varying concentrations of menthol were added individually to a VITROCELL® nebulizer and uniform aerosol clouds were made according to the manufacturer’s instructions. Control cells were exposed to aerosols created using 200 µL of 0.8% NaCl solutions containing 2 µL of 99% DMSO (equivalent to the amount of DMSO in the highest dose), which was prepared immediately prior to nebulization. Cells were exposed for 1.5 minutes to menthol aerosol nebulized in the cloud chamber, returned to the incubator for 4 h, then exposure was repeated a second time, after which they were placed in the incubator for 24 h.

**Menthol EC aerosol exposure at the ALI in a** **Cultex RFS compact exposure module**

A Cultex RFS compact exposure module (Cultex Laboratories GmbH, Hannover, Germany) was used to expose EpiAirway tissues to humidified zero air and EC aerosol using a vacuum pump that operates with a flow rate of 5 mL/min/insert. Six cell culture inserts were exposed at a time in each experimental run. Direct exposure of cells to EC aerosol was carried out using a custom designed (RTI International, North Carolina) EC smoking robot. The Cultex modular system consisted of the aerosol guiding module and the sampling module. The mouthpiece of the EC tank model was connected to a T connector which fits into the aerosol guiding module. For uniform distribution, the EC aerosol was diluted with humidified zero air (1 L/min) before being drawn into the Cultex® RFS compact exposure system. Puff conditions were as follows: 30 puffs with a puff volume of 55 mL, 4 sec puff duration and 60 sec inter-puff intervals. This puffing regimen was selected to deliver robust and consistent puff volumes. A programmable pump with a flow control application was used for dispensing aerosol. The aerosol guiding module (which is in the form of a T connector) was fitted tightly to the sampling module (Cultex RFS compact module) before exposure. The sampling module housed six culture inserts, which were separately supplied with medium. Exposure to humidified zero air was used as a control. Airflows were maintained and controlled by mass flow controller (Bronkhorst, Bethlehem, PA). Before exposure, the apical surfaces of the EpiAirway tissues were rinsed once with 500 µL PBS and then transferred to the exposure module containing 3 mL of maintenance medium/well. During each exposure, six tissue samples were exposed individually to EC aerosol diluted with humidified zero air or humidified zero air (control tissues). Control tissues were subject to the same procedures as test tissues, and a minimum of three tissue samples were included in each experiment. 24 h after exposure, medium was collected and stored to carry out cytotoxicity assays and ELISAs. EpiAirway tissue in the transwell inserts were used for the MTT assay and proteomics analysis.

**TRPM8 protein detection by immunohistochemistry**

BEAS-2B cells were cultured in chamber slides to approximately 75% confluence and then fixed with 4% paraformaldehyde in tris-buffered saline (TBS) for 10 minutes. Cells were washed three times with TBS containing 0.1% tween-20 (TBS/T), and non-specific binding was blocked using a solution of 10% donkey serum and 5% BSA in TBST. The cells were rinsed three times with TBS and incubated at 4^0^C for 18 h with a rabbit polyclonal IgG antibody (1:500) specific to human TRPM8 (Abcam, Cambridge, MA). Cells were washed and then treated for 1 h at room temperature with an Alexa-Fluor 488 conjugated donkey anti-rabbit IgG secondary antibody (Life Technologies, Carlsbad, CA) at a dilution of 1:400 in the blocking solution. Nuclei were counter-stained with 6-diamidino-2-phenylindole (DAPI) at 1:1000 dilution in TBS. Negative controls consisted of cells treated with secondary antibody alone. Images were collected using an inverted microscope (Nikon Ellipse) equipped with filters to visualize green fluorescent protein and DAPI. An integrated CMOS camera was used to collect images at 60X magnification. FITC and DAPI stained images were superimposed using ImageJ software. Immunoreactivity of TRPM8 was detected as green fluorescence.

**MTT assay**

Cell viability was measured by reduction of tetrazolium salt methyl thiazoyl tetrazolium (MTT) after 24 h of treatment in submerged culture and after EC aerosol exposure in the Cultex system. The MTT assay was performed 24 h after the second menthol aerosol exposure in the VITROCELL® ALI system.

For submerged culture with BEAS-2B cells, the MTT assay was performed as described in detail previously (Behar et al., 2018). 4,500 cells were seeded in 96-well plates. Cells were treated with menthol solution and aerosols diluted with culture medium. 24 h after treatment, MTT solution was added for 2 h, after which MTT medium was removed, and the purple formazan product was extracted by adding 200 µL of DMSO. 100 µL of cell extract from each treatment group were transferred to a clear 96-well plate in duplicates, and absorbance was read at 570 nm with a Synergy HTX Microplate Reader (Bio Tek, VT). Three independent experiments were performed and means and standard error of the mean (SEM) were used to produce concentration-response curves for each cell type.

For ALI exposures, 2 mL of MTT concentrate was prepared in 10 mL of BEGM medium just prior to use. Apical surfaces of the transwells were rinsed with PBS. 720 mL of BEGM medium with MTT was added to the basal layer of each transwell. Cells were incubated at 37^o^ C, 5% CO_2_ in an incubator for 2 h, after which MTT medium was removed, and the purple formazan product was extracted by adding 300 µL of DMSO to the apical layer. 100 µL of cell extract from each treatment group were transferred to a clear 96-well plate in duplicates, and absorbance was read at 570 nm with a Synergy HTX Microplate Reader (BioTek, VT). Each experiment was done three times, absorbance data were normalized to the incubator control, and the mean and standard error of the mean were determined.

**Live cell imaging of menthol-treated cells**

BEAS-2B cells were plated in 24-well plates at 8000 cells/well and incubated at 37^o^C for 24 h. Cells were treated with 0.02 mg/mL or 0.2 mg/mL of menthol solution or menthol aerosol fluid and incubated in a Nikon BioStation CT (Nikon, Melville, New York). Time-lapse images were captured every 2 h for 48 h. One set of cells was plated for 24 h (attached) before treatment and incubated in the BioStation, while the other set (attaching) was plated simultaneously with treatment. Menthol concentrations were chosen that did not produce any effect in the MTT assay. For each well, data were collected from five different fields. Images were segmented and analyzed using CL Quant software (DR Vision) to determine the rate of growth and morphology of the control and treated cells. To determine if cells that rounded up during incubation were dead, cells were treated with 0.4% trypan blue, and the percentage of dead cells was determined.

**Intracellular calcium imaging**

BEAS-2B cells (approximately 6000 cells/ well) were plated on 8-well Ibidi chamber slides (Ibidi, Munich, Germany) and left to attach for 24 h. Cells were transfected with GCaMP5 plasmid using DNA-In transfection reagent in Opti-MEM I Reduced Serum medium. 24 h after transfection, medium was removed and replaced with fresh BEGM medium, and the chamber slide with transfected cells was placed in a microscope stage top incubator chamber for live cell imaging using a Nikon Ellipse inverted microscope. Live cell videos of transfected cells were recorded. XY positions of each cell were registered and medium was replaced with medium containing 0.2 mg/mL menthol. Time-lapse videos were recorded before menthol treatment and 1, 2, 15 and 20 mins after addition of menthol. Inhibition of TRPM8 receptor was carried out using a TRPM8 receptor antagonist - N(4-tert-butylphenyl)-4-(3-chloropyridin-2-yl)piperazine-1-carboxamide (BCTC-10 µM), as previously reported (Sabnis et al 2008). Cells were treated with BCTC for 20 mins before and during menthol treatment. The change in fluorescence after menthol treatment for each time point with and without inhibitor was recorded using an Andor camera and analyzed using CL Quant software.

**Mitochondrial ROS measurement**

Mitochondrial ROS (superoxide) measurement was carried out in submerged cultures. BEAS-2B cells were grown on cover-glass 8 well chamber slides for 24 h, then treated with 0.2 mg/mL of menthol solution for 4 h. Media with menthol was removed, and cells were washed with PBS solution. Cells were then incubated with MitoSOX dye for 2 mins at 37^o^ C, after which they were washed and examined with a Nikon Eclipse inverted microscope. Images were collected using non-saturating exposure times. Controls were handled similarly but received no methanol treatment.

**Detection of mitochondrial protein oxidation using Mitotimer**

BEAS-2B cells were seeded (60,000 cells/insert) and allowed to attach for 24 h in transwell inserts. Cells were transfected with MitoTimer plasmid (Addgene, Cambridge, MA) using DNA-In transfecting reagent according to the manufacturer's instructions. Briefly, the cells were incubated with a mixture of 1 µg of plasmid DNA and 3 µl of DNA In (MTI-GlobalStem, MD) in Opti-MEM medium at 37^o^C. After 24 h, medium with the transfecting reagent was removed from the apical layer and fresh medium was added to the basal layer. The transfected cells were exposed to menthol aerosol that was generated using a nebulizer in a VITROCELL® cloud chamber. Menthol concentration was chosen that did not produce any effect in the MTT assay. Menthol aerosol was generated by adding 0.8 mg/mL menthol in 0.8% sodium chloride solution to the nebulizer. Each experiment included untreated control samples that remained in the incubator and controls that were exposed to only 0.8% sodium chloride (no menthol). 24 h after exposure, cells were fixed on inserts using 4% formaldehyde and the membranes were transferred to glass slides and mounted using Vectashield with DAPI (Vector Lab, Burlingame, CA). Fluorescence microscopy was performed using a Nikon Ellipse inverted microscope equipped with filters to visualize green (excitation/emission 488/518 nm) and red (excitation/emission 543/572 nm) channels. Images were collected from both channels using non-saturating exposures.

**Western blotting**

24 h after menthol aerosol exposure in the VITROCELL® cloud chamber, cells in transwells were lysed using RIPA buffer. To evaluate NF-κB activation, nuclear and cytoplasmic fractions of cell suspensions were isolated after treatment using the Nuclear/Cytosol Fractionation Kit (Thermo Fischer Scientific). Western blotting was performed as previously described (Nair et al., 2014). Briefly, lysates were collected and vortexed every 15 min for 1 h, centrifuged at 10,000 × g for 10 min at 4°C, and quantified using the Pierce BCA assay kit (Thermo Scientific, Waltham, Massachusetts). 20 µg of protein were used for loading each lane in gels. Lysates were mixed with Laemmli buffer (1:4), and proteins were separated using SDS gel electrophoresis (100 V for 2 h). The separated proteins were transferred to a PVDF membrane (BioRad, Carlsbad, CA) by wet electroblotting. The membrane was blocked with 5% milk in TBST buffer for 45 min and incubated overnight at 4^o^C with antibodies against TRPM8 (Abcam, Cambridge, MA), SOD2 (Cell Signaling Technology, Danvers, MA), and β-actin (Cell Signaling Technology, Danvers, MA). The membrane was washed for 30 min in TBST and incubated in secondary antibody. After 2 h, the membrane was developed using immunoblot reagent (BioRad, Hercules, California‎) in a ChemiDoc™ Imaging Systems (BioRad, Hercules, California‎).

**Quantification of cytokines (IL6 and IL8) in response to menthol treatment**

Conditioned medium from the basal layer of transwells was collected 24 h post-exposure and stored at -80^o^ C. Proinflammatory cytokine release was determined using IL-6 and IL-8 ELISA kits according to the manufacturer’s instructions (Invitrogen, Carlsbad, CA).

**siRNA Interference assay**

BEAS-2B cells were seeded in 12-well transwell inserts in serum free BEGM medium without antibiotics and allowed to attach for 24 h. Knockdown of TRPM8 along with iLamin as a control was carried out using siRNA and lipofectamine RNAiMAX according to the manufacturer's instructions. 24 h after transfection, medium was removed, and fresh medium was added to the basal layer. Transwells were exposed to menthol aerosol (0.8 mg/mL) in the VITROCELL® cloud chamber. 24 h after exposure, expression of proteins was analyzed using western blotting. siRNAs against TRPM8 and Lamin were prepared by Sigma-Aldrich.

**EC and chemicals**

An Innokin iTaste MVP 3.0 battery with variable voltage (V) and wattage (W) with fresh unused SMOK Pyrex Aro bottom coil tanks and a resistance of 1.8 ohms and a voltage of 3 V (4.7 watts) was used to generate EC aerosols using a custom-built smoking machine. Tanks were loaded with 2 mL of lab-made refill fluid and used in a manner that avoided dry puffing. Five initial puffs from each tank were used to prime the device before exposures to cells were made.

**Transepithelial electrical resistance (TEER) assay**

Barrier integrity of the EpiAirway epithelial tissue tight junctions was determined by measuring transepithelial electrical resistance (TEER) with an EVOM2 voltohmmeter and a 12 mm EndOhm electrode chamber (World Precision Instruments, Sarasota, Florida). Before TEER measurements, the apical surface of the tissues was rinsed three times with PBS, after which 500 µL of TEER buffer was added to the apical layer. Inserts were placed in a EVOM cup, and TEER readings were taken. The background resistance without the epithelial barrier present was subtracted from all measurements. The raw resistance (after background subtraction) was multiplied by 1.12 (surface area of culture insert) resulting in final values with units of X • cm^2^. TEER measurements following exposure are presented as the percentage of the pre-exposure value normalized to the control.

**Biological Assays for Cultex Exposed Tissues**

24 h after exposure, culture medium was collected from the basal side of each EpiAirway transwell, and aliquoted and stored at -80°C to carry out lactate dehydrogenase (LDH) cell death assays and IL-6 and IL-8 secretion analysis.

The LDH assay was performed using the Pierce LDH Cytotoxicity Assay Kit (Rockford, IL) for all samples, including negative and positive controls. For full kill (positive control) readings, 0.5% triton-X was added to the apical side of non-treated EpiAirway inserts for 2.5 h and placed in the incubator. Briefly, 50 µL of each sample and control, were pipetted in duplicate into a 96-well plate with 50 µL of Reaction Mixture. After a 30-minute incubation, 50 µL of Stop Solution were added. The absorbance was read across all samples at 490 nm and 680 nm; the latter was subtracted as background to give LDH activity which was computed as follows: % Cytotoxicity = (Sample activity – Clean Air Control activity) / (Full Kill Activity – Clean Air Control activity) x 100.

IL-6 and IL-8 concentrations were measured using enzyme-linked immunosorbent assays (ELISAs) as specified on the manufacturers protocol for each pro-inflammatory cytokine (Bender MedSystems, Vienna, Austria; Life Technologies, Carlsbad, CA, USA). Prior to running the assays, IL-6 samples were diluted 1:2 using culture medium and IL-8 samples were diluted 1:100 using the standard dilution buffer supplied in the kit. Four replicate EpiAirway samples were run for each treatment/control group.

**Protein Isolation and Proteome Processing of Cultex Exposed Tissues:**

Twenty-four hours after exposure in the Cultex, transwell membranes were removed by cutting, and the EpiAirway epithelial tissues were washed twice with PBS then lysed using RIPA buffer by vortexing for 1 min every 15 min for 45 min at 4^0^C. Lysates were centrifuged, and protein concentrations were determined using the BCA assay. 20 µg of protein were then subjected to SDS-polyacrylamide gel electrophoresis (PAGE), and the resulting gel was stained with Coomassie Brilliant Blue R-250 to visualize the separated proteins. The purpose of the gel was to ensure equal protein loading for each sample.

Lysates containing 150 µg of protein were precipitated with cold acetone (final concentration 80%) overnight at -20^o^C. The samples were centrifuged for 30 min at 14,000 rpm and the pellet was subjected to proteome analysis.

Protein pellets were treated with 10 µL of trypsin solution (0.1 mg/ml stock solution in 50 mM ammonium bicarbonate supplemented with 10% acetonitrile) (Roche Life Science) overnight at 37^o^C. The samples were placed on top of a vortex mixer for continuous agitation to keep pellets in suspension. After trypsin digestion, samples were centrifuged and supernatants were collected and dried down as pellets with a speedvac concentrator and re-dissolved in 20 µl 0.1% formic acid, which was the final sample then subjected to liquid chromatography (LC) / mass spectroscopy (MS) analysis.

A MudPIT approach was employed to analyze the trypsin-treated samples. A two-dimensional nanoAcquity ultra-pressure liquid chromatography (Waters, Milford, MA) and an Orbitrap Fusion MS (ThermoFisher Scientific, San Jose, CA) were configured to perform online 2D-nanoLC/MS/MS analysis. 2D-nanoLC was operated with a 2D-dilution method that is configured with nanoAcquity UPLC. The two mobile phases for the first-dimension LC fractionation were 20 mM ammonium formate (pH 10) and acetonitrile, respectively. Online fractionation was achieved by 5-minute elution off a NanoEase trap column (PN# 186003682, Waters) using a stepwise-increased concentration of acetonitrile. A total of five fractions were generated with 11%, 16%, 20%, 25%, and 50% of acetonitrile, respectively. A final flushing step used 80% acetonitrile to clean up the trap column. Each fraction was then analyzed online using a second-dimension LC gradient. The second dimension nano-UPLC method was described previously (Drakakaki et al., 2012).

The Orbitrap Fusion MS method was based on a data-dependent acquisition (DDA) survey. The acquisition time was set from 1-70 min. A Nano ESI source was used with the spray voltage at 2600V, sweep gas at 0, and ion transfer tube temperature at 275^o^C. An Orbitrap mass analyzer was used for the MS1 scan with resolution set at 60,000. MS mass range was 350-1800 m/z. The AGC target for each scan was set at 500,000 with maximal ion injection time set at 100 ms.

For the MS2 scan, the Orbitrap mass analyzer was used in an auto/normal mode with resolution set at 30,000. Only precursor ions with intensities of 50,000 or higher were selected for the MS2 scan. The sequence of individual MS2 scanning was from most-intense to least-intense precursor ions using a top-speed mode under time control of 4 sec. Higher energy CID (HCD) was used for fragmentation activation with 30% normalized activation energy. Quadrupole was used for precursor isolation with a 2 m/z isolation window. The MS2 mass range was set to auto/normal with the first mass set at 100 m/z. Maximal injection time was 100 ms with the AGC target set at 20,000. Ions were injected for all available parallelizable time. A 20-sec exclusion window was applied to all abundant ions to avoid repetitive MS2 scanning on the same precursor ions using 10 ppm error tolerance. Only charge states from 2 to 6 were allowed for MS2 scan, and undetermined charge states were not included. All MS2 spectra were recorded in the centroid mode.

The raw MS files were processed and analyzed using the Proteome Discoverer version 2.1 (ThermoFisher Scientific, San Jose, CA). The Sequest HT search engine was used to match all MS data to the human Uniprot protein database supplemented with common contaminant proteins such as keratins. The search parameters were the following: trypsin with 2 missed cleavage, minimal peptide length for six amino acids, MS1 mass tolerance 20 ppm, MS2 mass tolerance 0.05 Da, Gln→pyro-Glu (N-term Q), oxidation (M), N-terminal acetylation as variable modification. Only proteins with a 1% false discovery rate (FDR) cut-off were considered in the final result.

**Statistical Analysis**

Analysis of submerged and VITROCELL® protocols: Absorbance data for the MTT and ELISAs were normalized to the untreated control (submerged cultures) or to the incubator control, and the means and standard errors of the means were determined using GraphPad Prism. For the MTT assays, the inhibitory concentrations at 50% (IC_50_) and 70% (IC_70_) values were computed with Prism software (GraphPad, San Diego, California, USA) using the log inhibitor versus normalized response-variable slope with the top and bottom constraints set to 100% and 0%, respectively.

For experiments on calcium influx with and without TRPM8 inhibitor, fluorescence was quantified relative to the control (without menthol) using CL-Quant software. The statistical significance for these data was calculated using a 2-way analysis of variance ANOVA with Bonferroni’s multiple comparison test. All time points in each treatment group (menthol without inhibitor and menthol with inhibitor) were compared.

A two-tailed t-test was used to analyze the mitochondrial ROS measurement (MitoSOX) and SOD2 expression using western blots. MitoTimer mitochondrial protein oxidation experiments, MTT and ELISA were analyzed using a one-way ANOVA. When significance was found, menthol exposed groups were compared to the control using Dunnett’s post hoc test.

Analysis of Proteomics Data: For all cell-based assays (MTT, LDH, ELISAs and TEER), an ANOVA with Dunnett’s post hoc test was carried out to compare the Clean Air Control (CA CN) to the other groups (PG control, and Menthol + PG). For the proteomics datasets, statistical analysis was done in a manner described in Statistics Analysis Supplemental to obtain adjusted p-values and fold changes for each significant protein. Proteins with known significant p-values and without a given fold-change were assigned 100x and 0.01x fold changes for upregulation and downregulation, respectively.

To identify groups of proteins sharing common pathways, we submitted a list of significantly altered proteins to DAVID (Huang et al., 2009) and the Ingenuity® Pathway Analysis (IPA®) omics analysis tool to discover networks and pathways of interest.

Data Access: The mass spectroscopy proteomics data (raw files) will be deposited into the ProteomeXchange Consortium.

**Work Cited**

Behar, R.Z., Davis, B., Wang, Y., Bahl, V., Lin, S., Talbot, P., (2014). Identification of toxicants in cinnamon-flavored electronic cigarette refill fluids. Toxicol. Vitr. 2014 March;(28): 198–208. DOI: https://doi.org/10.1016/j.tiv.2013.10.006

Behar, R.Z., Hua, M., Talbot, P., (2015). Puffing topography and nicotine intake of electronic cigarette users. [PLoS One.](https://www.ncbi.nlm.nih.gov/pubmed/25664463) 2015 Feb 9;10(2):e0117222. DOI:[10.1371/journal.pone.0117222](https://doi.org/10.1371/journal.pone.0117222)

Behar, R.Z., Luo, W., Lin, S.C., Wang, Y., Valle, J., Pankow, J.F., Talbot, P., (2016). Distribution , quantification and toxicity of cinnamaldehyde in electronic cigarette refill fluids and aerosols. Tob Control. 2016 Sep 15;25: ii94–ii102. DOI: 10.1136/tobaccocontrol-2016-053224

Behar, R.Z., Wang, Y., Talbot, P., (2017). Comparing the cytotoxicity of electronic cigarette fluids, aerosols and solvents. [Tob Control.](https://www.ncbi.nlm.nih.gov/pubmed/28596276) 2018 May;27(3):325-333. DOI: [10.1136/tobaccocontrol-2016-053472](https://doi.org/10.1136/tobaccocontrol-2016-053472)

Drakakaki, G., Van De Ven, W., Pan, S., Miao, Y., Wang, J., Keinath, N.F., Weatherly, B., Jiang, L., Schumacher, K., Hicks, G., Raikhel, N., (2012). Isolation and proteomic analysis of the SYP61 compartment reveal its role in exocytic trafficking in Arabidopsis. Cell Res. 2012 Feb;22: 413–424. DOI: 10.1038/cr.2011.129

Huang, D.W., Sherman, B.T., Lempicki, R.A., (2009). Systematic and integrative analysis of large gene lists using DAVID bioinformatics resources. Nat. Protoc. 2009;4(1): 44–57. DOI: 10.1038/nprot.2008.211

Nair, V., Sreevalsan, S., Basha, R., Abdelrahlm, M., Abudayyeh, A., Hoffman, A.R., Safe, S., (2014). Mechanism of metformin-dependent inhibition of mammalian target of rapamycin (mTOR) and Ras activity in pancreatic cancer. J. Biol. Chem. 2014 Oct 3;289(40): 27692–27701. DOI: 10.1074/jbc.M114.592576

Sabnis, A.S., Shadid, M., Yost, G.S., Reilly, C.A., (2008). Human lung epithelial cells express a functional cold-sensing TRPM8 variant. [Am J Respir Cell Mol Biol.](https://www.ncbi.nlm.nih.gov/pubmed/18458237) 2008 Oct;39(4):466-74. DOI: [10.1165/rcmb.2007-0440OC](https://doi.org/10.1165/rcmb.2007-0440OC)
