## Supplementary material for "Menthol in Electronic Cigarettes: A Contributor to Respiratory Disease?": Statistics Analysis Supplemental

**In-House Statistical Model for Determination of Significant Proteins and Fold Changes in Proteomic Data**

**Department of Applied Statistics**

**University of California, Riverside**

1. CA vs PG vs CAD
   1.1. Pre-Deal with Dataset

Dataset are collected with the count of number of proteins, say x. We did the transformation x → log2(x + 0.5), to make sure that our dataset are approximately normally distributed. Further,

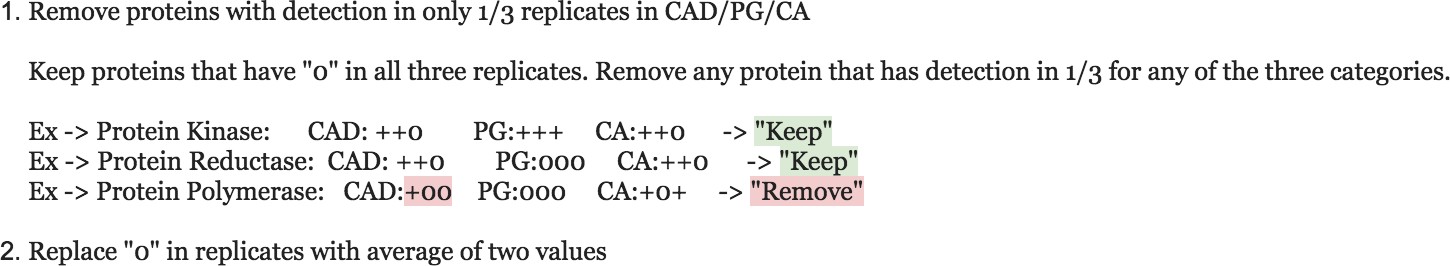

#### 1.1. Introduction

In this section, we want to test the treatment efficacy of CAD on protein expression when compared to control cells (CA CN). Since the treatments were delivered using a propylene glycol (PG) solvent, we must also test how the solvent altered protein expression together with CA and CAD. In other words, the null hypothesis would be

H_0_: CA = PG = CAD*,*

it is equivalent to the union of 3 null hypotheses

H^(1)^: CA = PG	vs	H^(1)^: CA $\neq$ PG

H^(2)^: CA = CAD	vs	H^(2)^: CA $\neq$ CAD

H^(3)^: PG = CAD	vs	H^(3)^: PG $\neq$ CAD

#### 1.3. Two-Layer Multiple Comparison Test

##### 1.3.1. Layer I

In Layer I, we focused on a single protein. We have 3 groups: 1 = CA, 2 = PG, 3 = CAD. In each group, we have 3 replicates. Let µi be the average protein expression for group i, i = 1, 2, 3, our parameters of interests are therefore {µi −µj}, 1≤j<i≤3. We will perform **Tukey’s Honest Significant Difference (HSD) Test** to build a simutaneous confidence interval of {µi − µj}, 1≤j<i≤3, with 90% coverage probability (significant level α = 0.1). Check Appendix A for the theoretical method of Tukey’s HSD Test.

A protein would be identified as significant if the corresponding simutaneous CI for {µi − µj}1≤j<i≤3 fails to contain (0, 0, 0). More specific, if the lower bound of CI for µi − µj is larger than 0, we would say that the average protein expression of group i is larger than group j (or µi > µj); on the other hand, if the upper bound of CI for µi − µj is smaller than 0, we would say that the average protein expression of group i is smaller than group j (or µi < µj).

##### 1.3.2. Layer II - PFER (Per Family Error Rate) Control

In Layer I, for a single protein *k*, we built a simutaneous CI with 1 *− α* coverage probability. Let *V_k_* be the number of cases where group *i* and group *j* are tested to be significantly different when they are actually not, 1 *≤ j < i ≤* 3 for protein *k* (*V_k_ ≥* 1 *⇔* protein *k* would

be falsely discovered as significant), thus,

P(*V_k_ ≥* 1) = *α*

In Layer II, we want to identify all the proteins that are considered to be significant. Let V* be the number of proteins that are tested to be significant when they are actually not, we have

$E[V*]=E[\overset{N}{\underset{i=1}{\sum}}I_{(V_{k}\geq1)}]=\overset{N}{\underset{k=1}{\sum}}P(V_{k}\geq1)=N\alpha$,

where *N* is the total number of proteins.

### 2. CA vs PG vs Menthol

Similar.

### 3. CA vs INC

#### 3.1. Two-Layer Single Comparison Test

The only difference is in Layer I.

#### 3.2. Layer I

For a single protein, we want to compare between 2 groups, 1 = CA, 2 = INC. For each group, we have 3 replicates. Since we only have 2 groups to compare, it turns out to be the Simple T-Test. Check Appendix B for the theoretical method of Simple T-Test.

### A. Tukey’s HSD Test

Let $y_{ij}$ be the observation of group i for j-th replicate, i = 1, ..., m; j = 1, ..., n. There are m

groups, each group has total n replicates (in our case, n = m = 3). Let

$$\overline{y}_{i}=\frac{1}{n}\overset{n}{\underset{j=1}{\sum}}y_{ij},i=1,...,m$$

$$s_{i}^{2}=\frac{1}{n-1}\overset{n}{\underset{j=1}{\sum}}(y_{ij}-\overline{y}_{i})^{2},i=1,...,m$$

$$S_{p}^{2}=\frac{1}{m}\overset{m}{\underset{i=1}{\sum}}S_{i}^{2}$$

$\overline{y}_{i}$ is the estimate of *µ_i_*;

$S_{p}^{2}$ is called the pooled estimated of variance.

Let Q stands for the Studentized Range Distribution, with number of groups = m and degrees of freedom = m(n-1), set significant level = α, then

*T* = *Q*(1 *− α, m, m*(*n −* 1))

*HSD* = $TS_{p}\sqrt{\frac{1}{n}}$

*T* is the upper *α* point for *Q* distribution.

And the simutaneous CI for $\{\mu_{i}-\mu_{j}{\}}_{i\neq j}$ would be

P(*µ_i_ − µ_j_ ∈* [*y*¯ *_i_ − y*¯ *_j_ ± HSD*]*, i* $\neq$ *j*) = 1 *− α.*

### B. Simple T-Test

Set m = 2, which means there are only 2 groups (in our case, they are INC and CA).

Let t stands for the Student t Distribution, with degrees of freedom = m(n-1),

$$H=t(1-\frac{\alpha}{2},m(n-1))$$

$$LSD=HS_{p}\sqrt{\frac{2}{n}}$$

H is the upper $\frac{\alpha}{2}$ point for t distribution. And the CI for *µ*_1_ *− µ*_2_ would be

P(*µ*_1_ *− µ*_2_ *∈* [*y* ¯ _1_ *− y*¯ _2_ *± LSD*]) = 1 *− α.*
